# Aging limits the reduction of reactive muscle activity to perturbations but spares the retrieval of context-specific patterns with practice

**DOI:** 10.64898/2026.11.11.766012

**Authors:** Shuqi Liu, Andrea M Weinstein, Gelsy Torres-Oviedo

**Author notes:** **Corresponding Author:** Dr. Gelsy Torres-Oviedo.

## Abstract

Moving smoothly through a changing environment requires immediate reactive responses to maintain balance at an environmental transition, and learned, context-specific responses to efficiently meet the new environmental demands. People use both the reactive and context-specific responses to adapt to new environment and to re-adapt more rapidly on subsequent re-encounters. While re-adaptation slows with older age, the underlying muscular mechanisms are unknown. We hypothesized that practice would reduce reactive recruitment and increase context-specific recruitment and that both changes would be smaller in older adults. To test this, younger and older adults practiced split-belt walking (one belt faster than the other) over three days. We recorded surface electromyography from 10 muscles per leg and used linear regression to decompose each session’s activity into a reactive component (template from naïve, early exposure) and a context-specific component (template from practiced activity). Mixed-effects models showed that, with practice, older adults decreased reactive recruitment less than younger adults, particularly in distal muscles, with no strong evidence for an age difference in context-specific recruitment. In sum, with practice, aging limits the reduction of reactive responses to a novel walking context but spares the retrieval of context-specific muscle patterns, suggesting that mobility rehabilitation for older adults could target reactive mechanisms.

## Introduction

Walking safely through changing environments requires humans to react to unfamiliar conditions and to retrieve previously learned walking patterns when a familiar condition is encountered again. For example, when walking on an icy sidewalk for the first time, people rely on reactive responses to avoid falling while developing a context-specific walking pattern that meets the task demands, such as a wider stance and/or slower speed. After repeated encounters, people react less and instead retrieve this context-specific pattern, transitioning to the icy sidewalk more efficiently. Faster re-adaptation upon re-encountering a previously experienced condition is known as “locomotor savings”^1–4^. Locomotor savings could therefore arise from a reduction of reactive responses, from more efficient retrieval of context-specific patterns, or from both. Prior work has shown that aging impairs locomotor savings^5,6^, but these studies quantified savings with a single behavioral variable, which reflects both processes together and cannot distinguish between them. Thus, it remains unknown whether aging limits the reduction of reactive responses, the retrieval of context-specific patterns, or both.

The reactive and context-specific responses can be studied by decomposing muscle activity into reactive and context-specific components, whose recruitment explains how walking movements adapt to environmental changes ^7,8^. Upon an environmental transition, reactive muscle activity is engaged as a balance response^7^ to the perturbation and then reduced as interaction with the new environment continues ^9–11^. Meanwhile, a context-specific muscle activity pattern is gradually forged ^7^ and could represent a learned response to improve walking efficiency in the new environment ^12,13^. This dual process of reactive and context-specific responses mirrors the stabilizing feedback controller to avoid falling and a memory controller, respectively, in the hierarchical framework for locomotor adaptation proposed by previous work^14^. Furthermore, muscle activity reflects neural drive from cortical and subcortical areas and can be assessed non-invasively ^15,16^, as opposed to other metrics that could occur with passive interactions with the environment (kinematics^17^) or are highly invasive (implanted electrodes in the brain). Therefore, characterizing how reactive and context-specific muscle activity patterns are impacted by aging and by practice over multiple days can provide insight into the neural control underlying the formation and retrieval of walking memories.

We hypothesized that the recruitment of the reactive muscle activity pattern would decrease and the recruitment of the context-specific pattern would increase with practice in both age groups, but that these changes would be smaller in older adults, contributing to slower re-adaptation when re-encountering an environmental change. To test this, younger and older adults experienced multiple sessions of a well-established paradigm ^1,2,4^, the split-belt treadmill, where each leg moves at a different speed, to assess changes in the reactive and context-specific muscle activity patterns after practice. We expected that older adults would continue to react more to the split-belt perturbation and recruit less of the context-specific muscle activity pattern than younger adults even after practice. Finally, we asked whether this age-related difference would also be detectable using a simpler measure based on overall muscle activity magnitude, which has been shown to be sensitive to age-related decline in motor learning ^8^.

## Results

Twenty-one older adults and 21 sex-matched younger adults completed a split-belt walking protocol as previously reported ^5^ to evaluate the changes in the reactive and context-specific muscle patterns after practice. Surface electromyography (EMG) was recorded from 10 muscles per leg spanning the hip, knee, and ankle (full list and abbreviations in Methods). Participant demographics are shown in Table 1. Both groups had comparable self-selected overground walking speeds (Table 1, t = 0.97, *p* = 0.34), measured as the average speed while walking back and forth on a straight 9.2m walkway for 150 strides (Figure 1a, baseline overground). Functional gait assessment (FGA) was administered in older adults (Table 1) to measure dynamic balance and functional mobility ^18^. The older adult cohort was physically healthy as indicated by the observation that 18 out of 21 older adults scored above the fall-risk cutoff (22/30, ^19–21)^. Older adults were screened for no signs of cognitive impairment, indicated by a Montreal Cognitive Assessment (MoCA ^22^) score greater than or equal to 24. Full inclusion criteria are given in the Methods.

**Figure 1.**
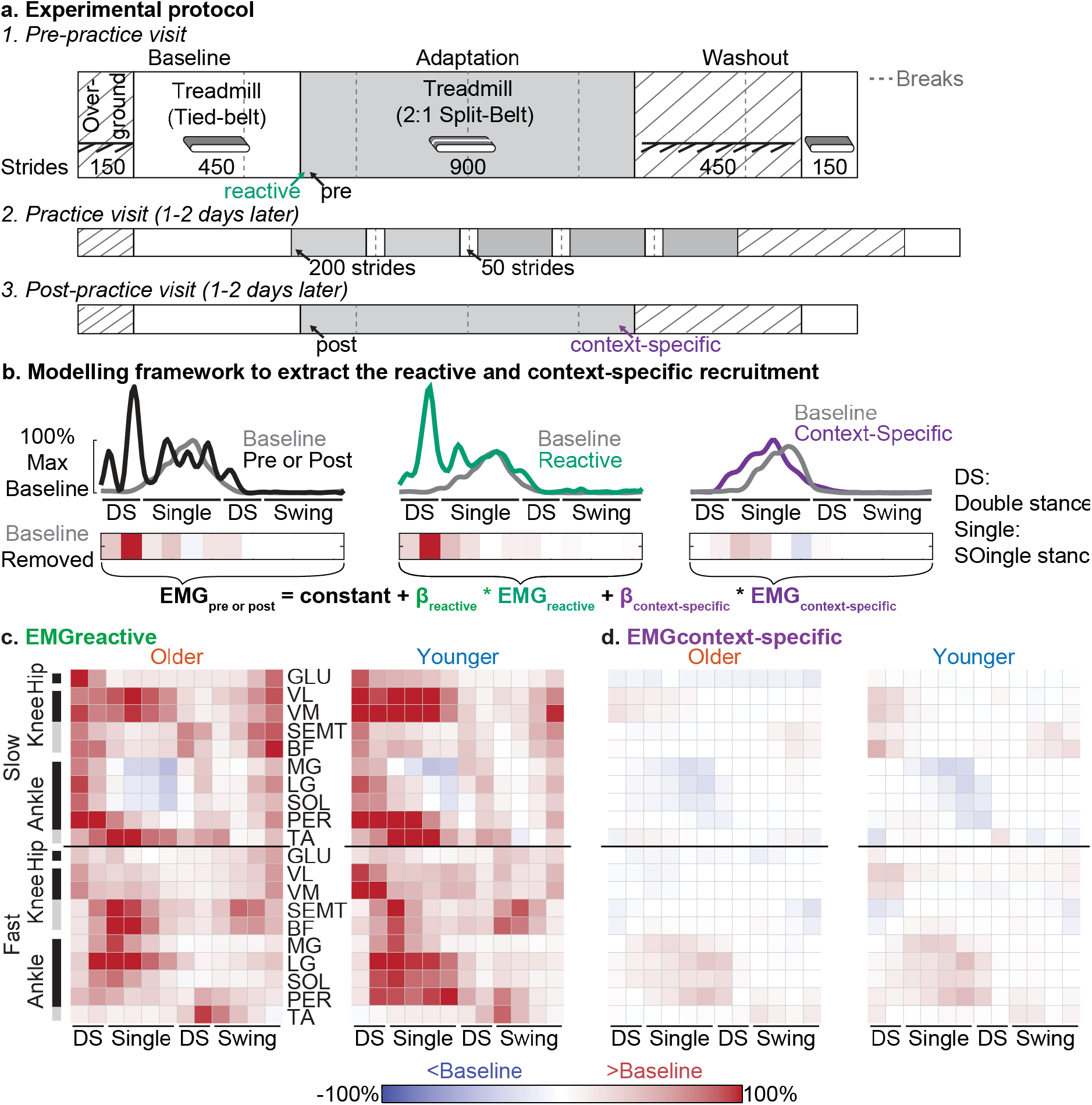
Experimental paradigm and electromyographic (EMG) computational framework. **(a)** Schematics of the split-belt treadmill walking paradigm to evaluate changes in reactive and context-specific muscle activity patterns after practice. Participants completed three visits on separate days: pre-practice, practice, and post-practice with each visit 1 or 2 days apart. In the pre-practice session, participants learned a new walking pattern on a split-belt treadmill where two legs are driven at different speeds. Patched blocks represent self-paced overground walking, white blocks represent tied-belt treadmill walking, grey blocks represent split-belt walking, and vertical dashed grey lines represent breaks. We analyzed EMG activity during early adaptation at the pre- and post-practice sessions. **(b)** Linear regression modeling framework used to extract reactive and context-specific muscle activity patterns. EMG activity for each muscle was binned into 12 sub-intervals per gait cycle, amplitude-normalized such that 100% corresponds to peak baseline activity, and baseline-subtracted (positive values represent activity above baseline; negative values represent activity below baseline). The recruitment of reactive and context-specific patterns per muscle was quantified via regression coefficients ( β_reactive_and β_context–specific_) by regressing early adaptation EMG activity onto two regressor templates: EMG_reactive_(measured immediately after split-belt onset in the pre-practice visit) and EMG_context–specific_ (measured at late adaptation in the post-practice visit). **(c)** Measured reactive muscle activity for each group. Each heatmap shows group-mean activity from proximal to distal muscles, ordered from the fast (f) to the slow (s) leg. Red indicates activity above baseline; orange indicates activity below baseline. Although the analysis was performed at the individual level, group-mean activity is shown here for visualization only, to convey the general characteristics of the reactive and context-specific muscle activity patterns and to allow comparison with previous studies ^7,8,23^. **(d)** Measured group-mean context-specific muscle activity. Notice that colors are lighter overall than the reactive activity in (c), indicating that context-specific activity after practice was closer to baseline.

**Table 1.** Participant characteristics.

| **Variables** | **Older Adults**<br>**Mean (SD) or n (%)** | **Younger Adults**<br>**Mean (SD) or n (%)** |
|---|---|---|
| Sample size | 21 | 21 |
| Age (years) | 71.85 ± 3.14 | 25.05 ± 5.20 |
| Female, n (%) | 11 (52.38%) | 11 (52.38%) |
| Race, n (%) |  |  |
| White | 19 (90.48%) | 16 (76.19%) |
| Black or African American | 1 (4.76%) | 1 (4.76%) |
| More than one race | 1 (4.76%) | 0 |
| Asian | 0 | 4 (19.05%) |
| Body mass index (BMI) | 26.82 ± 4.55 | 25.05 ± 5.20 |
| Right leg dominant, n (%) | 21 (100%) | 20 (95.24%) |
| Self-selected overground walking speed (m/s) | 0.96 ± 0.17 | 1.01 ± 0.14 |
| Functional Gait Assessment (FGA) score (max 30) | 24.71 ± 2.43 | not administered |

**Table 1.**
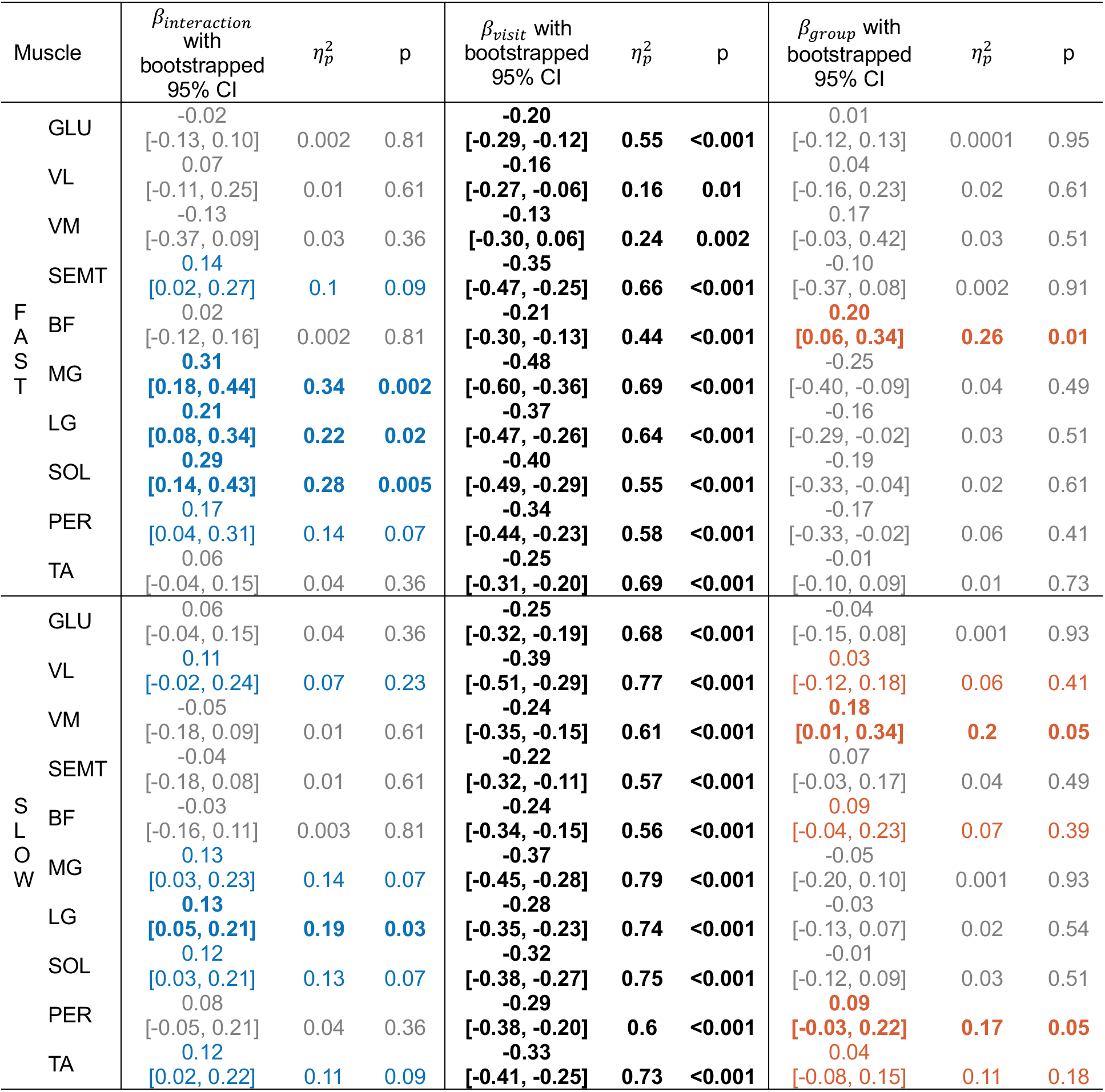
β_reactive_ second-level linear mixed effect model results. The reported β_interaction_, β_visit_, β_7roup_ and 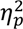 are the model estimate derived from the observed β_reactive_ data extracted from the first-level model. The 95% confidence interval was obtained via bootstrapping. P-values were computed by permutation testing and adjusted for multiple comparisons across 20 muscles using the Benjamini–Hochberg false discovery rate (FDR) procedure ^31^. Effects remaining significant after correction are formatted in bold. Observed medium or large interaction effect sizes (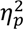 = 0.06 ^32^) are shown in blue when the change in older adults is smaller than younger adults (matching expectation) and in orange vice versa, consistent with the color schemes in Figures 3, 4, and 5. Small or negligible effects are in grey. Observed visit effects that are medium or large are shown in black. Observed medium or large group effects are shown in orange when recruitment is greater in older adults (expected) and in blue when recruitment is smaller in older adults.

| Muscle |  | β<sub>interaction</sub> with bootstrapped 95% CI | η<sub>p</sub><sup>2</sup> | p | β<sub>visit</sub> with bootstrapped 95% CI | η<sub>p</sub><sup>2</sup> | p | β<sub>group</sub> with bootstrapped 95% CI | η<sub>p</sub><sup>2</sup> | p |
|---|---|---|---|---|---|---|---|---|---|---|
| F<br>A<br>S<br>T | GLU | -0.02<br>[-0.13, 0.10] | 0.002 | 0.81 | **-0.20**<br>**[-0.29, -0.12]** | **0.55** | **<0.001** | 0.01<br>[-0.12, 0.13] | 0.0001 | 0.95 |
|  | VL | 0.07<br>[-0.11, 0.25] | 0.01 | 0.61 | **-0.16**<br>**[-0.27, -0.06]** | **0.16** | **0.01** | 0.04<br>[-0.16, 0.23] | 0.02 | 0.61 |
|  | VM | -0.13<br>[-0.37, 0.09] | 0.03 | 0.36 | **-0.13**<br>**[-0.30, 0.06]** | **0.24** | **0.002** | 0.17<br>[-0.03, 0.42] | 0.03 | 0.51 |
|  | SEMT | 0.14<br>[0.02, 0.27] | 0.1 | 0.09 | **-0.35**<br>**[-0.47, -0.25]** | **0.66** | **<0.001** | -0.10<br>[-0.37, 0.08] | 0.002 | 0.91 |
|  | BF | 0.02<br>[-0.12, 0.16] | 0.002 | 0.81 | **-0.21**<br>**[-0.30, -0.13]** | **0.44** | **<0.001** | **0.20**<br>**[0.06, 0.34]** | **0.26** | **0.01** |
|  | MG | **0.31**<br>**[0.18, 0.44]** | **0.34** | **0.002** | **-0.48**<br>**[-0.60, -0.36]** | **0.69** | **<0.001** | -0.25<br>[-0.40, -0.09] | 0.04 | 0.49 |
|  | LG | **0.21**<br>**[0.08, 0.34]** | **0.22** | **0.02** | **-0.37**<br>**[-0.47, -0.26]** | **0.64** | **<0.001** | -0.16<br>[-0.29, -0.02] | 0.03 | 0.51 |
|  | SOL | **0.29**<br>**[0.14, 0.43]** | **0.28** | **0.005** | **-0.40**<br>**[-0.49, -0.29]** | **0.55** | **<0.001** | -0.19<br>[-0.33, -0.04] | 0.02 | 0.61 |
|  | PER | 0.17<br>[0.04, 0.31] | 0.14 | 0.07 | **-0.34**<br>**[-0.44, -0.23]** | **0.58** | **<0.001** | -0.17<br>[-0.33, -0.02] | 0.06 | 0.41 |
|  | TA | 0.06<br>[-0.04, 0.15] | 0.04 | 0.36 | **-0.25**<br>**[-0.31, -0.20]** | **0.69** | **<0.001** | -0.01<br>[-0.10, 0.09] | 0.01 | 0.73 |
| S<br>L<br>O<br>W | GLU | 0.06<br>[-0.04, 0.15] | 0.04 | 0.36 | **-0.25**<br>**[-0.32, -0.19]** | **0.68** | **<0.001** | -0.04<br>[-0.15, 0.08] | 0.001 | 0.93 |
|  | VL | 0.11<br>[-0.02, 0.24] | 0.07 | 0.23 | **-0.39**<br>**[-0.51, -0.29]** | **0.77** | **<0.001** | 0.03<br>[-0.12, 0.18] | 0.06 | 0.41 |
|  | VM | -0.05<br>[-0.18, 0.09] | 0.01 | 0.61 | **-0.24**<br>**[-0.35, -0.15]** | **0.61** | **<0.001** | **0.18**<br>**[0.01, 0.34]** | **0.2** | **0.05** |
|  | SEMT | -0.04<br>[-0.18, 0.08] | 0.01 | 0.61 | **-0.22**<br>**[-0.32, -0.11]** | **0.57** | **<0.001** | 0.07<br>[-0.03, 0.17] | 0.04 | 0.49 |
|  | BF | -0.03<br>[-0.16, 0.11] | 0.003 | 0.81 | **-0.24**<br>**[-0.34, -0.15]** | **0.56** | **<0.001** | 0.09<br>[-0.04, 0.23] | 0.07 | 0.39 |
|  | MG | 0.13<br>[0.03, 0.23] | 0.14 | 0.07 | **-0.37**<br>**[-0.45, -0.28]** | **0.79** | **<0.001** | -0.05<br>[-0.20, 0.10] | 0.001 | 0.93 |
|  | LG | **0.13**<br>**[0.05, 0.21]** | **0.19** | **0.03** | **-0.28**<br>**[-0.35, -0.23]** | **0.74** | **<0.001** | -0.03<br>[-0.13, 0.07] | 0.02 | 0.54 |
|  | SOL | 0.12<br>[0.03, 0.21] | 0.13 | 0.07 | **-0.32**<br>**[-0.38, -0.27]** | **0.75** | **<0.001** | -0.01<br>[-0.12, 0.09] | 0.03 | 0.51 |
|  | PER | 0.08<br>[-0.05, 0.21] | 0.04 | 0.36 | **-0.29**<br>**[-0.38, -0.20]** | **0.6** | **<0.001** | **0.09**<br>**[-0.03, 0.22]** | **0.17** | **0.05** |
|  | TA | 0.12<br>[0.02, 0.22] | 0.11 | 0.09 | **-0.33**<br>**[-0.41, -0.25]** | **0.73** | **<0.001** | 0.04<br>[-0.08, 0.15] | 0.11 | 0.18 |

### The naïve split-belt muscle activity resembles reactive balance responses, and the expert muscle activity represents the context-specific responses

Figure 1a shows the experimental protocol. Faster adaptation when re-encountering a perturbation can be quantified by comparing early adaptation behavior pre- and post-practice. To understand the muscular mechanisms, we decomposed the early adaptation EMG activity in both pre- and post-practice visits into two components: a reactive and a context-specific component (Figure 1b), using linear regression following the previously established method ^7^. Note that all muscle activity in the regression equation (both dependent and independent variables) was recorded, and the linear regression simply quantified the reactive and context-specific activity contributions and how the contributions change over practice and between age groups.

Figures 1c and 1d show the group-mean reactive and context-specific muscle activity patterns used as regressors. The reactive pattern (Figure 1c) was measured by the naïve response to first exposure to the split-belt treadmill (mean early-adaptation EMG over strides 2–5) and is consistent with reactive balance responses reported previously ^8,23–28^. The context-specific pattern was measured by the expert responses during late adaptation in the post-practice visit. Qualitatively, the context-specific pattern (Figure 1d) is consistent with prior reports ^8,12,29,30^ and shows lower overall activity than the reactive pattern (Figure 1d, most colors are close to white).

### In calf muscles, older adults recruited less reactive muscle activity pattern than younger adults pre-practice in the fast leg, but more than younger adults post-practice in the slow leg

In our main analysis, we expected that practice would result in a smaller decrease in the recruitment of the reactive muscle activity pattern in older than younger adults, which would be detected as a group-by-visit interaction. Examples of how the recruitment of the reactive muscle activity pattern (*β_reactive_*) changes during adaptation were shown in Figure 2a and 2b for a fast medial gastrocnemius (MG) and slow lateral gastrocnemius (LG), respectively. The reactive muscle activity pattern was engaged immediately and decreased gradually with adaptation. Both groups recruited less reactive pattern during early adaptation at the post-practice compared to the pre-practice visit, but the change was smaller in older than younger adults, corresponding to a group-by-visit interaction.

**Figure 2.**
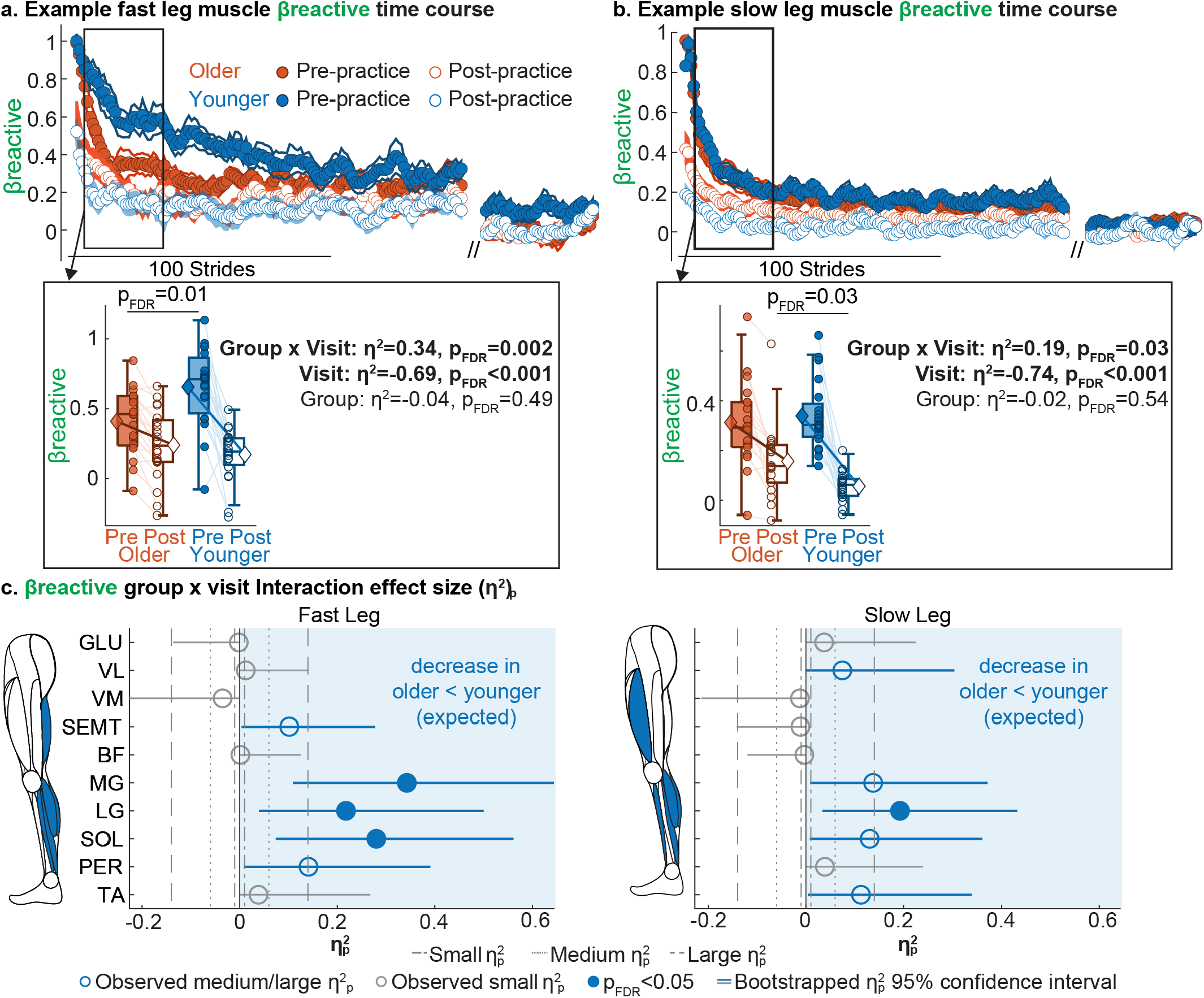
Time courses of the reactive muscle activity pattern recruitment with illustration of the group and visit effect analysis in example fast (a) and slow (b) leg muscles and summary of the group x visit interaction effect. **(a)** Time course of the recruitment of the reactive activity (β_reactive_) for fast-leg medial gastrocnemius (MG) during adaptation. Older adults are shown in orange, younger adults in blue; pre-practice and post-practice data are in closed and open circles, respectively. Each dot represents the average of 5 consecutive strides, and shaded area represents the standard error. β_reactive_ decreased during adaptation in both groups and was smaller at the post-practice. Black box indicates the stride window where the data was analyzed to determine the visit, group and interaction effect shown in the box below. Insert: boxplot visualizing β_reactive_ from individual participants. Each dot represents one participant. Thin lines connect data from the same participant. Group means are shown as diamonds connected by thick lines. The slopes were different between groups, corresponding to a significant interaction effect. Both slopes were negative corresponding to a significant negative visit effect (β_reactive_ decreased post practice). There was no overall difference between orange and blue, corresponding to a non-significant group effect. **(b)** Time course of β_reactive_ for slow-leg lateral gastrocnemius (LG). **(c)** Interaction effect size 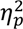 for β_reactive_ per muscle. A schematic leg colors muscles with medium or large effects in blue for effects in the expected direction where older adults showed smaller changes than younger adults. The background and the observed effects that were medium or large in the expected direction are colored blue; medium or large effects in the opposite direction (older adults changing more than younger adults) are colored orange; small or negligible effects are shown in grey. A schematic leg provides a visual for the anatomical locations of the muscles with the same coloring scheme. Dots represent the observed effect size; the line represents the 95% confidence interval from bootstrapping. Filled dots indicate a significant interaction after Benjamini– Hochberg false discovery rate (FDR) ^31^ correction. Dot-dashed, dotted, and dashed lines mark reference thresholds for small (0.01), medium (0.06), and large (≥0.14) effect sizes ^32^.

The interaction effect in *β_reactive_* was quantified for all muscles in Figure 2c. We found significant group-by-visit interactions with large effect sizes matching our expectation (Table 1, Figure 2c, filled blue) in fast-leg calf muscles (MG, LG, soleus, SOL) and slow LG. All effects were robust to outlier removal (i.e., effect sizes remained large after outlier removal). Fast-leg calf results were specific to the adaptation condition, i.e., the interaction was larger in adaptation than in the baseline condition (MG, *p* = 0.01; LG, *p* = 0.01; SOL, *p* = 0.047; supplementary Table S1, Figure S3). Post-hoc analysis revealed that the two groups recruited the reactive pattern differently at the pre-practice visit in the fast-leg muscles, but at the post-practice visit in the slow-leg muscles (supplementary Figure S1). Specifically, older adults showed significantly smaller recruitment of reactive pattern than younger adults at the pre-practice visit in fast MG (*p* = 0.01 after Benjamini–Hochberg false discovery rate (FDR) ^31^ *correction*; Figure 2a insert) and SOL (*p* = 0.03), with a trending difference in fast LG (*p* = 0.07), while no group difference was observed post-practice. On the other hand, in slow LG, the recruitment was similar between group pre-practice, but significantly higher in older than younger adults post-practice (Figure 2b insert, *p* = 0.03).

Trending (0.05 < *p* < 0.09) medium to large interactions in the same direction (Figure 2c open blue circles) were observed in other slow-leg calf muscles (MG, peroneus, PER), tibialis anterior (TA), and fast leg semitendinosus (SEMT), although the effect size in fast SEMT became small after outlier removal. A non-significant medium to large interaction was also observed in slow leg vastus lateralis (VL). All *β_reactive_* data are shown in supplementary Figure S1.

Taken together, our results suggest that older adults exhibit smaller practice-dependent reduction of the reactive muscle activity pattern than younger adults in distal muscles, but for different reasons in each leg. In the fast leg, older adults started less reactive but reached similar post-visit recruitment like younger adults, whereas in the slow leg older adults started similar to younger adults but showed limited reduction of the reactive pattern post-practice.

### Older adults recruited less context-specific muscle activity pattern after practice than younger adults in fast tibialis anterior (TA)

In the main analysis for the recruitment of the context-specific muscle activity pattern, we expected a smaller practice-dependent increase in *β_context_*_–*specific*_ in older than in younger adults, which would be detected as a group-by-visit interaction. Examples of how the recruitment of context-specific muscle activity pattern ( *β_context_*_–*specific*_) changes during adaptation were shown in Figure 3a and 3b for fast and slow TA, respectively. The context-specific pattern was recruited gradually during adaptation. Both groups recruited more context-specific muscle activity pattern during early adaptation at post-than at pre-practice visit. The difference between visits was smaller in older than younger adults in fast TA, corresponding to a significant interaction in fast TA.

**Figure 3.**
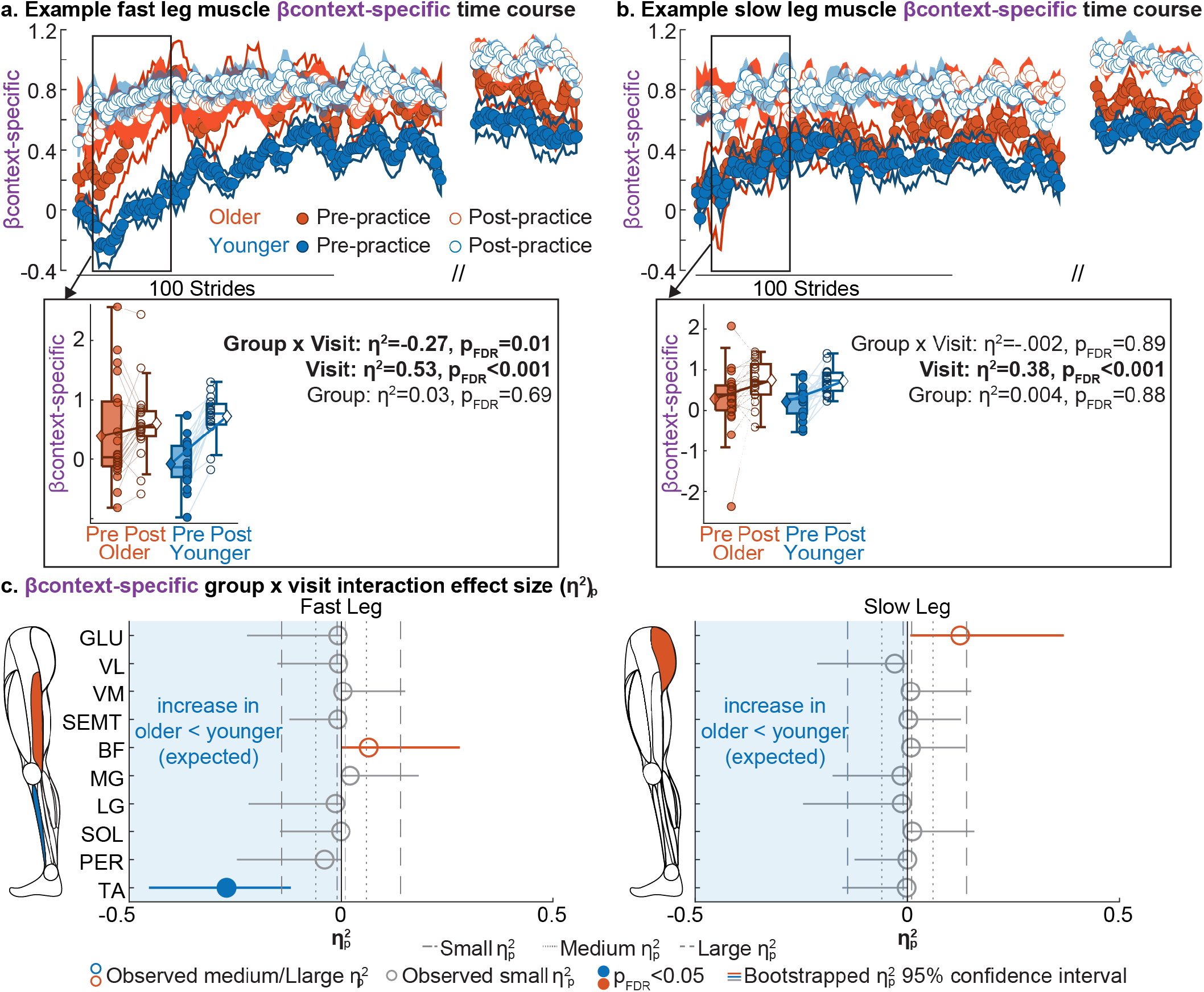
Context-specific muscle activity pattern recruitment time courses with illustration of the group and visit effect analysis in example fast (a) and slow (b) leg muscles and summary of the group x visit interaction effect. **(a)** Time course of the recruitment of the context-specific activity (β_context–specific_) for fast-leg tibialis anterior (TA) during adaptation. Older adults data are shown in orange, younger adults in blue; pre-practice and post-practice data are shown as closed and open circles, respectively. Each dot represents the average of 5 consecutive strides, and shaded area represents the standard error. β_context–specific_increased during adaptation and was retrieved at the post-practice visit (higher β_context–specific_). Black box indicates the stride window where the data was analyzed to determine the visit, group and interaction effect shown in the box below. Insert: Boxplot visualizing β_context–specific_ with one dot per participant. Thin lines connect data from the same participant. Group means are in diamonds connected by thick lines. The slopes were different between groups, corresponding to a significant interaction effect. Both slopes were positive corresponding to a significant positive visit effect ( β_context–specific_increased post practice). There was no overall difference between orange and blue, corresponding to a non-significant group effect. **(b)** Time course of β_context–specific_ for slow-leg TA. **(c)** Interaction effect size 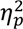 for β_context–specific_ per muscle. A schematic leg colors muscles with medium or large effects in blue for effects in the expected direction where older adults showed smaller increase than younger adults in β_context–specific_ from practice, and in orange vice versa. The blue background marks the expected direction. Observed effects that were medium or large in the expected direction are colored blue; medium or large effects in the opposite direction are colored orange; small or negligible effects are shown in grey. Dots represent the observed effect size; the line represents the 95% confidence interval from bootstrapping. Filled dots indicate a significant interaction after Benjamini–Hochberg false discovery rate (FDR) ^31^ correction. Dot-dashed, dotted, and dashed lines mark reference thresholds for small (0.01), medium (0.06), and large (≥0.14) effect sizes ^32^.

The interaction effect in *β_context_*_–*specific*_ was quantified for all muscles in Figure 3c. A significant group-by-visit interaction with large effect size matching our expectation was observed in fast TA (Table 2, Figure 3c, filled blue). This effect was robust to outlier removal and was specific to the adaptation condition (*p* = 0.002). Post-hoc analysis revealed no significant difference between groups at either visit. Non-significant interaction in the opposite direction with medium to large sizes were observed in fast biceps femoris (BF), slow gluteus medius (GLU), but both effects were sensitive to outlier removal, i.e., both effect sizes were small after outlier removal. All *β_context_*_–*specific*_ data are shown in supplementary Fig S2.

**Table 2.**
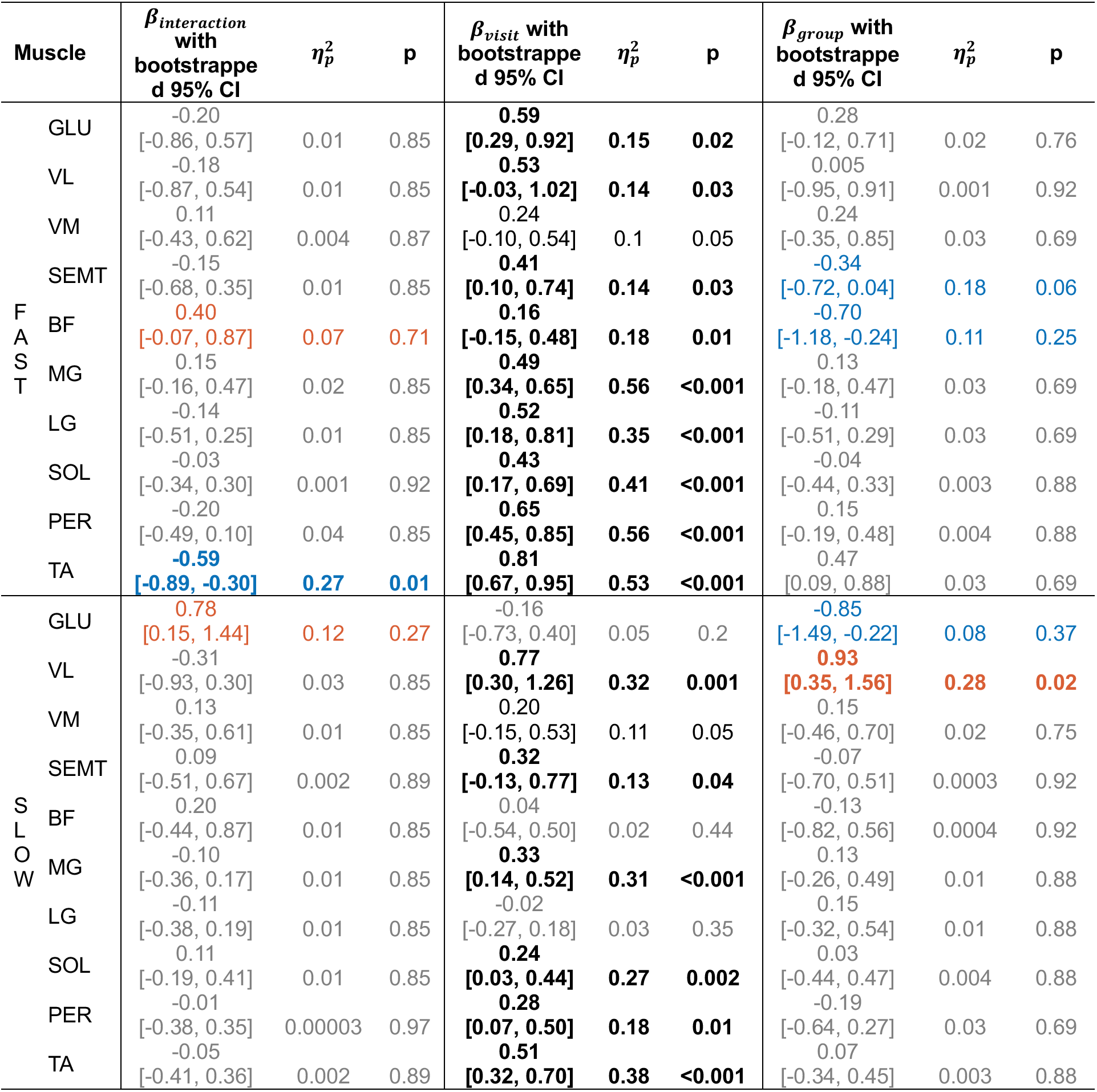
β_context–specific_ second-level linear mixed effect model results. The reported β_interaction_, β_visit_, β_7roup_ and 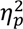 are the model estimate derived from the observed β_context–specific_ data extracted from the first-level model. The 95% confidence interval was obtained via bootstrapping. P-values were computed by permutation testing and adjusted for multiple comparisons across 20 muscles using the Benjamini–Hochberg false discovery rate (FDR) procedure ^31^. Effects remaining significant after correction are formatted in bold. Observed interaction effect that are medium or large (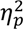 >= 0.06 ^32^) are shown in blue when the change in older adults is smaller than that in younger adults (matching expectation), and in orange vice versa, consistent with the color conventions in Figure 3, 4, and 5. Small or negligible effects are in grey. Observed medium or large visit effects are shown in black. Observed medium or large group effects are shown in orange when recruitment is greater in older adults and in blue when recruitment is smaller in older adults (expected).

| Muscle |  | β<sub>interaction</sub> with bootstrapped 95% CI | η<sub>p</sub><sup>2</sup> | p | β<sub>visit</sub> with bootstrapped 95% CI | η<sub>p</sub><sup>2</sup> | p | β<sub>group</sub> with bootstrapped 95% CI | η<sub>p</sub><sup>2</sup> | p |
|---|---|---|---|---|---|---|---|---|---|---|
| FAST | GLU | -0.20<br>[-0.86, 0.57] | 0.01 | 0.85 | **0.59<br>[0.29, 0.92]** | **0.15** | **0.02** | 0.28<br>[-0.12, 0.71] | 0.02 | 0.76 |
|  | VL | -0.18<br>[-0.87, 0.54] | 0.01 | 0.85 | **0.53<br>[-0.03, 1.02]** | **0.14** | **0.03** | 0.005<br>[-0.95, 0.91] | 0.001 | 0.92 |
|  | VM | 0.11<br>[-0.43, 0.62] | 0.004 | 0.87 | 0.24<br>[-0.10, 0.54] | 0.1 | 0.05 | 0.24<br>[-0.35, 0.85] | 0.03 | 0.69 |
|  | SEMT | -0.15<br>[-0.68, 0.35] | 0.01 | 0.85 | **0.41<br>[0.10, 0.74]** | **0.14** | **0.03** | -0.34<br>[-0.72, 0.04] | 0.18 | 0.06 |
|  | BF | 0.40<br>[-0.07, 0.87] | 0.07 | 0.71 | **0.16<br>[-0.15, 0.48]** | **0.18** | **0.01** | -0.70<br>[-1.18, -0.24] | 0.11 | 0.25 |
|  | MG | 0.15<br>[-0.16, 0.47] | 0.02 | 0.85 | **0.49<br>[0.34, 0.65]** | **0.56** | **<0.001** | 0.13<br>[-0.18, 0.47] | 0.03 | 0.69 |
|  | LG | -0.14<br>[-0.51, 0.25] | 0.01 | 0.85 | **0.52<br>[0.18, 0.81]** | **0.35** | **<0.001** | -0.11<br>[-0.51, 0.29] | 0.03 | 0.69 |
|  | SOL | -0.03<br>[-0.34, 0.30] | 0.001 | 0.92 | **0.43<br>[0.17, 0.69]** | **0.41** | **<0.001** | -0.04<br>[-0.44, 0.33] | 0.003 | 0.88 |
|  | PER | -0.20<br>[-0.49, 0.10] | 0.04 | 0.85 | **0.65<br>[0.45, 0.85]** | **0.56** | **<0.001** | 0.15<br>[-0.19, 0.48] | 0.004 | 0.88 |
|  | TA | **-0.59<br>[-0.89, -0.30]** | **0.27** | **0.01** | **0.81<br>[0.67, 0.95]** | **0.53** | **<0.001** | 0.47<br>[0.09, 0.88] | 0.03 | 0.69 |
| SLOW | GLU | 0.78<br>[0.15, 1.44] | 0.12 | 0.27 | -0.16<br>[-0.73, 0.40] | 0.05 | 0.2 | -0.85<br>[-1.49, -0.22] | 0.08 | 0.37 |
|  | VL | -0.31<br>[-0.93, 0.30] | 0.03 | 0.85 | **0.77<br>[0.30, 1.26]** | **0.32** | **0.001** | **0.93<br>[0.35, 1.56]** | **0.28** | **0.02** |
|  | VM | 0.13<br>[-0.35, 0.61] | 0.01 | 0.85 | 0.20<br>[-0.15, 0.53] | 0.11 | 0.05 | 0.15<br>[-0.46, 0.70] | 0.02 | 0.75 |
|  | SEMT | 0.09<br>[-0.51, 0.67] | 0.002 | 0.89 | **0.32<br>[-0.13, 0.77]** | **0.13** | **0.04** | -0.07<br>[-0.70, 0.51] | 0.0003 | 0.92 |
|  | BF | 0.20<br>[-0.44, 0.87] | 0.01 | 0.85 | 0.04<br>[-0.54, 0.50] | 0.02 | 0.44 | -0.13<br>[-0.82, 0.56] | 0.0004 | 0.92 |
|  | MG | -0.10<br>[-0.36, 0.17] | 0.01 | 0.85 | **0.33<br>[0.14, 0.52]** | **0.31** | **<0.001** | 0.13<br>[-0.26, 0.49] | 0.01 | 0.88 |
|  | LG | -0.11<br>[-0.38, 0.19] | 0.01 | 0.85 | -0.02<br>[-0.27, 0.18] | 0.03 | 0.35 | 0.15<br>[-0.32, 0.54] | 0.01 | 0.88 |
|  | SOL | 0.11<br>[-0.19, 0.41] | 0.01 | 0.85 | **0.24<br>[0.03, 0.44]** | **0.27** | **0.002** | 0.03<br>[-0.44, 0.47] | 0.004 | 0.88 |
|  | PER | -0.01<br>[-0.38, 0.35] | 0.00003 | 0.97 | **0.28<br>[0.07, 0.50]** | **0.18** | **0.01** | -0.19<br>[-0.64, 0.27] | 0.03 | 0.69 |
|  | TA | -0.05<br>[-0.41, 0.36] | 0.002 | 0.89 | **0.51<br>[0.32, 0.70]** | **0.38** | **<0.001** | 0.07<br>[-0.34, 0.45] | 0.003 | 0.88 |

Taken together, the age-by-visit interaction in the development of context-specific muscle activity pattern was only observed in fast TA where older adults showed less practice-induced increase.

### The recruitment of reactive and context-specific muscle activity patterns could help explain unexpected changes in EMG magnitude

To determine whether a conventional metric ^8,12,29^ would capture these age and practice related changes, we analyzed overall EMG magnitude. We expected older adults to show a smaller practice-related decrease in EMG magnitude than younger adults. Non-significant interactions with medium effect size in the expected direction were seen in fast-leg MG, SOL, slow-leg VL, vastus medialis (VM), and TA (supplementary Figure S5a, S6, Table S2, S3). These decreases in magnitude could be related to reduced recruitment of the reactive muscle activity pattern. Correspondingly, most of these muscles (fast-leg MG, SOL, slow-leg VL, TA) also showed interactions in *β_reactive_* in the same direction. An interaction contrary to the expectation was observed in fast-leg TA in EMG magnitude, and the interaction was related to younger adults showed a surprising increase in EMG magnitude after practice (supplementary figure S5a, S6). The increase in magnitude could reflect increased recruitment of the context-specific muscle activity pattern, and consistently, fast TA is also the only muscle with a significant interaction in *β_context_*_–*specific*_ with younger adults showing a significantly larger increase in *β_context_*_–*specific*_ from practice than older adults. In sum, results from EMG magnitude were largely consistent with those from *β_reactive_* and *β_context_*_–*specific*_, but information from the decomposed processes provided more details in aiding the interpretation of seemingly surprising findings from EMG magnitude.

### Practice reduced the recruitment of the reactive muscle activity pattern and increased the recruitment of the context-specific pattern

To better understand the impact of practice, we examined the main effect of visit as a secondary analysis. We expected that the recruitment of the reactive muscle activity pattern would decrease while the recruitment of the context-specific pattern would increase after practice in both groups, both of which would be detected as a significant visit effect.

As expected, in *β_reactive_*, a significant visit effect was observed in all muscles (Figure 4a; Table 1) where practice robustly reduced the recruitment of the reactive muscle activity pattern (negative slopes between visit within group as seen in example Figure 2a, 2b inserts). The visit effect was robust to outlier removal in all muscles. The effect was also specific to the adaptation condition in all muscles except fast-leg VL and VM (supplementary Figure S3, Table S1), suggesting that the *β_reactive_* reduction reflects adaptation to the experimental task rather than mere familiarity with the lab environment.

**Figure 4.**
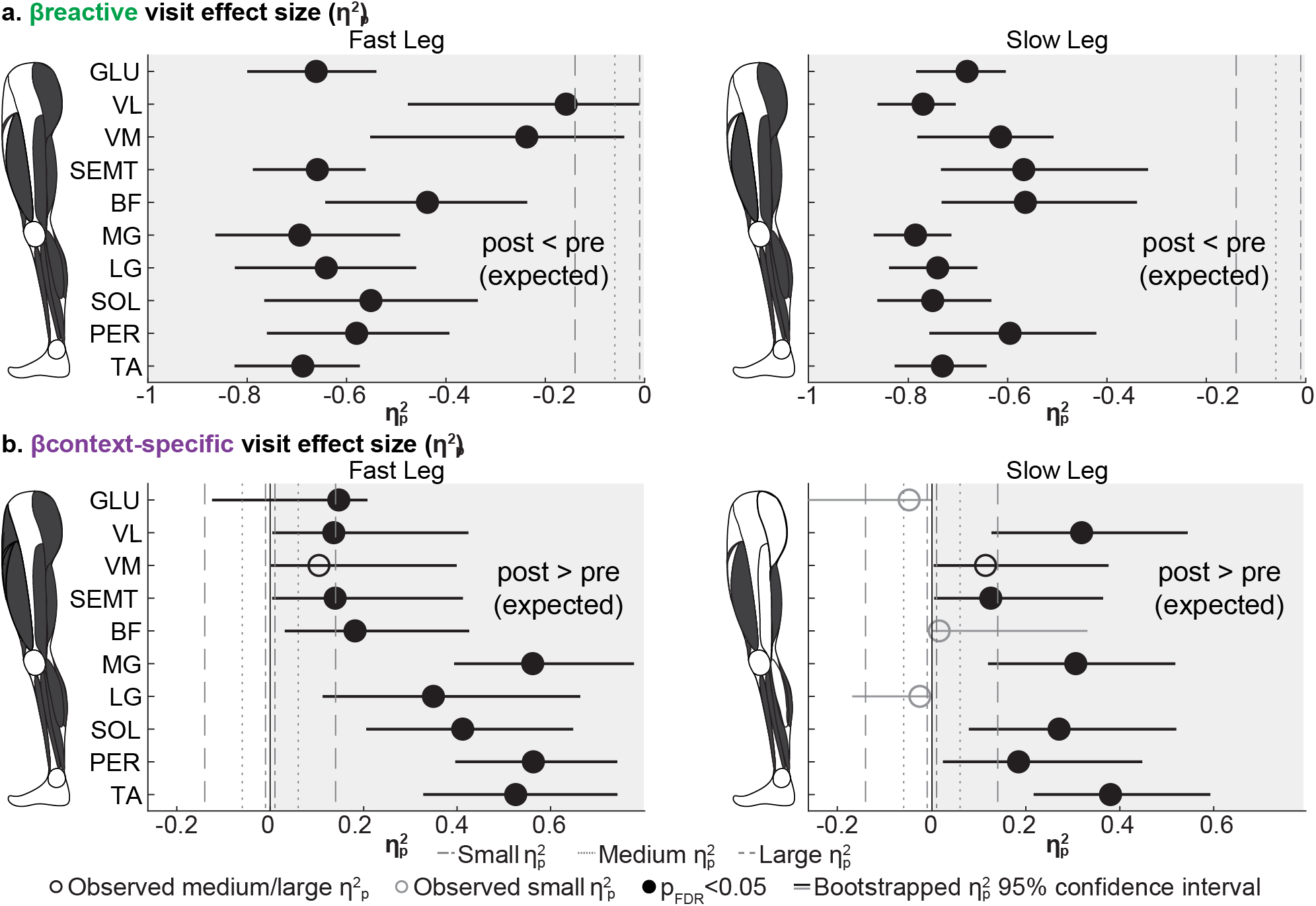
Main effect of visit in the recruitment of reactive (a) and context-specific (b) muscle activity. **(a)** Overview of the visit effect size 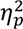 for reactive recruitment (β_reactive_) per muscle. A schematic leg colors the muscles with medium or large effects in dark grey. Each dot represents the observed effect size; the line represents the 95% confidence interval from bootstrapping. Filled dots indicate a significant effect after Benjamini–Hochberg false discovery rate (FDR) correction. The grey background marks the expected direction where the reactive recruitment decreased after practice. Observed effects that were medium or large in the expected direction are shown in black. Small or negligible effects are shown in grey. Dot-dashed, dotted, and dashed lines mark reference thresholds for small (0.01), medium (0.06), and large (≥0.14) effect sizes ^32^. **(b)** Overview of the visit effect size 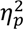 for context-specific recruitment (β_context–specific_) per muscle. Note that the expected direction is reversed: the context-specific recruitment was expected to increase after practice.

In *β_context_*_–*specific*_, significant visit effects with medium to large effect sizes matching our expectation were observed in all fast-leg muscles except VM and majority of the slow-leg muscles (VL, VM, SEMT, MG, SOL, PER, and TA; Figure 4b; Table 2), all showing higher context-specific recruitment post-practice (positive slopes within groups as seen in example Figure 3a, 3b inserts). All effects were robust to outlier removal. The visit effect was specific to the adaptation condition in fast GLU, all fast distal muscles (MG, LG, SOL, PER, and TA), slow SEMT, and slow TA (supplementary Figure S4, Table S1). Non-significant and medium to large effects were also observed in bilateral VM.

Lastly, the visit effect observed in EMG magnitude matched the results from *β_reactive_* except in fast-leg TA, where EMG magnitude increased after practice (supplementary Figure S5b, S6, Table S2, S3).

Taken together, these results indicate that both age groups adapted faster when re-encountering a perturbation via a similar muscular mechanisms: by reducing the use of the reactive muscle activity pattern after practice in all muscles and increasing the recruitment of the context-specific muscle activity pattern in the majority of the muscles. The reactive reduction was adaptation-specific in nearly all muscles, but the context-specific increase was adaptation-specific in only some, mostly fast-leg muscles, suggesting the increased recruitment of context-specific muscle activity pattern is not fully attributable to split-belt adaptation, particularly in the slow leg.

### Group differences were uncommon and distributed across different muscle groups and legs

To determine if the two age groups differed overall, we also examined the main effect of group as a secondary analysis. We expected that older adults would recruit more reactive and less context-specific muscle activity pattern than younger adults across visits.

In *β_reactive_*, fast-leg BF, slow-leg VM and PER showed a significant group effect matching our expectation where older adults recruited more reactive pattern than younger adults (Figure 5a, Table 1). These effects were robust to outlier removal, but none was specific to the adaptation condition (supplementary Figure S3, Table S1). Three additional slow-leg muscles spanning distinct anatomical groups (VL, BF, and TA) showed effects in the same direction (older > younger) with medium-to-large effect sizes, but none reached significance. The effect sizes in slow-leg VL and BF also became small after outlier removal.

**Figure 5.**
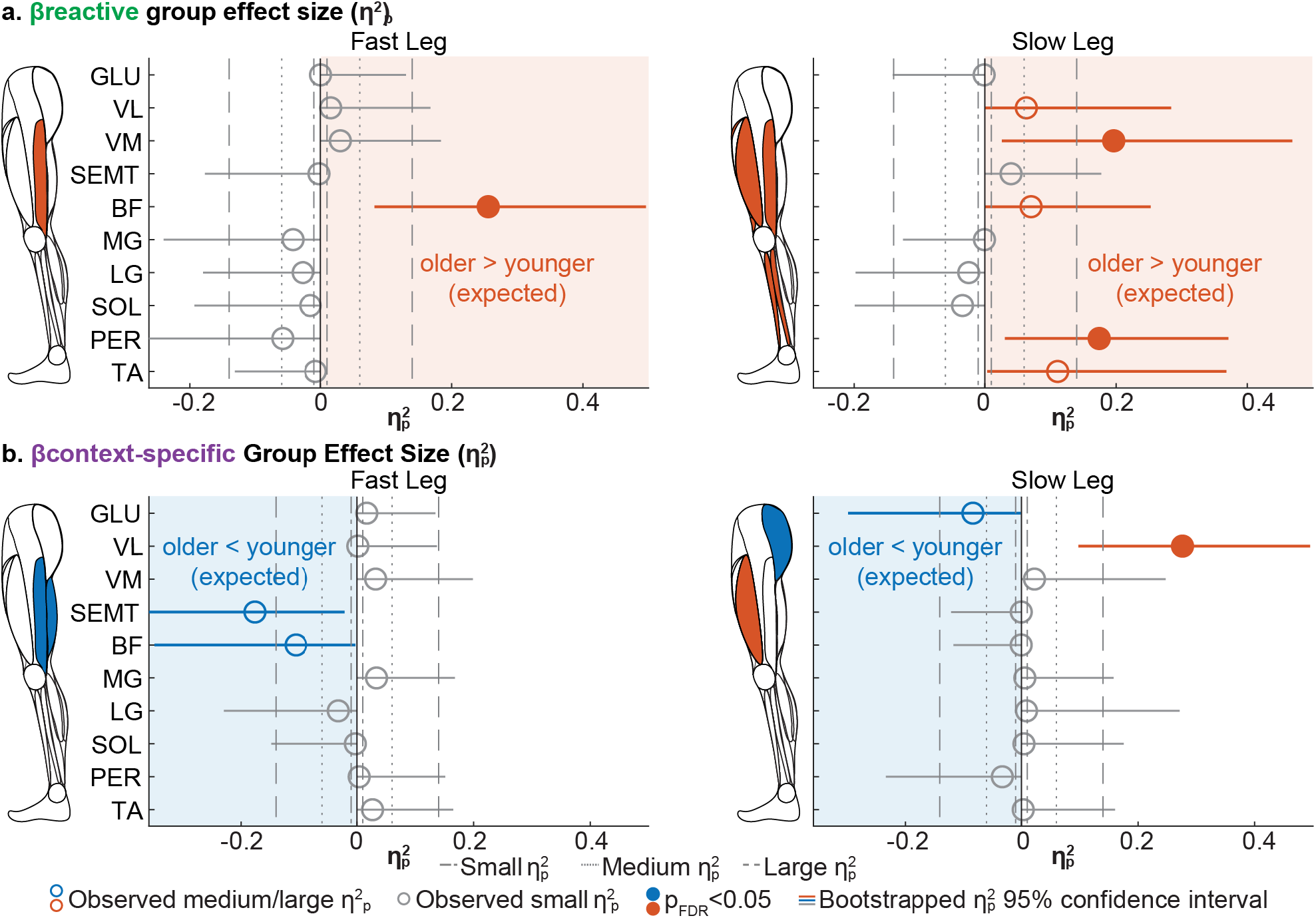
Main effect of group in the recruitment of reactive (a) and context-specific (b) muscle activity patterns. **(a)** Overview of the group effect size 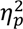 for reactive recruitment β_reactive_ per muscle. A schematic leg colors the muscles with medium or large effects in orange when the recruitment was greater in older adults, and in blue when the recruitment was smaller in older adults. Each dot represents the observed effect size; the line represents the 95% confidence interval from bootstrapping. Filled dots indicate a significant interaction after Benjamini–Hochberg false discovery rate (FDR) correction. The orange background marks the expected direction, in which older adults recruit more reactive pattern than younger adults. Observed effects that were medium or large in the expected direction (older > younger) are colored orange; medium or large effects in the opposite direction (older adults < younger adults) are colored blue; small or negligible effects are colored grey. Dot-dashed, dotted, and dashed lines mark reference thresholds for small (0.01), medium (0.06), and large (≥0.14) effect sizes ^32^. **(b)** Overview of the group effect size 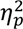 for context-specific recruitment (β_context–specific_) per muscle. Display conventions as in (a), except that the expected direction is reversed: older adults were expected to show less context-specific recruitment than younger adults, and therefore the background marks the expected (older < younger) side.

In *β_context_*_–*specific*_, slow-leg VL showed a significant group effect in the direction opposite than expected (Figure 5b, Table 2), where older adults recruited more context-specific pattern than younger adults. This effect was robust to outlier removal and specific to the adaptation condition (*p* = 0.03; supplementary Figure S4, Table S1). In contrast, three non-significant, medium or large effect in the expected direction was observed in fast-leg SEMT, BF and slow-leg GLU where older adults recruited less context-specific pattern than younger adults.

Lastly, EMG magnitude showed mixed results (supplementary Figure S5c, Table S2, S3). Several muscles showed larger magnitude in older than younger adults (bilateral SEMT, fast-leg BF, slow-leg MG, LG, TA), whereas fast-leg VL, VM and PER showed smaller magnitude in older than younger adults. EMG magnitude results largely diverged from *β_reactive_* and *β_context_*_–*specific*_.

Taken together, group effect was limited and observed in muscles across different legs and muscle groups. Specifically, the expected higher reactive recruitment in older adults was observed in fast-leg hamstring, slow-leg quadriceps, and slow-leg ankle plantar flexor (PER). While in the context-specific pattern, expected lower recruitment in older adults was only observed in fast-leg hamstrings, slow-leg hip; instead, an unexpected higher recruitment in older than younger adults was observed in slow-leg VL. Group effect in EMG magnitude was not consistently aligned with either decomposed metric.

## Discussion

We showed that older adults did not reduce the recruitment of the reactive muscle activity pattern as well as younger adults after practice, particularly in the distal muscles, whereas both age groups showed similar changes in the recruitment of the context-specific pattern. These results suggest that the age difference in the muscle control of faster re-adaptation when re-encountering a perturbation is associated mainly with a reduced ability to downregulate the reactive balance responses to a perturbation, even after practice, while the ability to form and recall context-specific motor memories is retained. In secondary analyses, we demonstrated that practice produced a robust reduction in reactive recruitment across all muscles and an increase in context-specific recruitment in most muscles, suggesting that both groups underwent similar muscular changes after practice but to different extents. Group main effects were uncommon and scattered across muscles and legs, with no strong evidence of an overall between-group difference in the recruitment of either pattern. Finally, electromyography (EMG) magnitude was largely inconsistent with the reactive or context-specific recruitment except for the visit effect, suggesting that the reactive and context-specific decomposition captures information distinct from EMG magnitude.

Age-by-visit interactions were largely confined to the recruitment of the reactive muscle activity pattern, with little evidence of interaction in context-specific recruitment. One explanation is that these two processes rely on different neural structures. The reactive process has been linked to cerebral contributions ^33^, while the context-specific process, resembling error-driven motor adaptation, may be more cerebellum-dependent ^34,35^. Aging is associated with widespread structural and functional decline in the brain ^36,37^, but emerging evidence suggests cerebellar function remains relatively resilient ^38–41^. Cerebral decline could thus impair practice-dependent adjustment of the reactive pattern, while preserved cerebellar function could support the formation of the context-specific pattern, consistent with prior work positing that internal-model recalibration does not deteriorate with aging ^41^. It also aligns with reports that older adults reach steady-state (late adaptation) behavior similar to younger adults in kinematics ^2,5,42–46^ and muscle activity magnitude ^8^, while differing in early adaptation ^5,43,45,47–49^.

The reactive recruitment interactions were concentrated in the distal leg muscles. This aligns with previous reports that proximal (upper-leg) muscles are less impacted by asymmetric walking conditions ^12,50^, whereas distal muscles stabilize the ankle and prepare for foot landing and placement ^12,30,51,52^. The reduced modulation of distal muscle activity in older adults could reflect a differential impact of aging on distal versus proximal muscles. Specifically, there is evidence that corticospinal excitability of distal muscles is reduced with age, whereas that of proximal muscles is not ^53^. The distal muscles could be more impacted by aging because they have longer axons, which are more susceptible to age-related changes such as sensory loss^54^ and decreased metabolism ^55^. Consistent with this, an age-related difference in balance strategy has been reported, in which younger adults preferentially use an ankle strategy, recruiting distal muscles first, whereas older adults use a hip strategy, recruiting proximal muscles first to restore balance^56–58^. If this preference extends to locomotor adaptation paradigms such as the split-belt walking task used here, younger adults may have relied more on ankle muscles at split-belt onset and refined this control with practice. This preferential impact of aging on distal muscles could explain the smaller change in the recruitment of reactive muscle activity in older than in younger adults.

The reactive recruitment interaction differed between the two legs. The slow leg behaved as expected with both age groups starting with similar reactive recruitment, and older adults showed smaller reduction of the recruitment compared to younger adults after practice. In the fast leg, by contrast, older adults showed a rapid reduction in reactive recruitment before practice and reached a similar recruitment level to younger adults after practice. This rapid reduction in older adults should not be interpreted simply as better tuning of the reactive responses. Instead, we suspect it reflects a poorer characterization of the reactive muscle activity pattern to the perturbation in the fast leg of older adults. This interpretation is supported by the goodness of fit (R², Supplementary Fig. S7) of the first level model from which the recruitment coefficient (*β_reactive_*) was extracted: it was significantly lower in older than in younger adults at the pre-practice visit for most fast distal muscles (MG, LG, SOL, and PER), whereas it was similar between groups at both visits for the slow leg. Thus, the rapid reduction in reactive recruitment in the fast distal muscles of older adults may indicate that these participants continued to adjust their muscle activity to counteract the perturbation early on, so that the epoch we chose to represent the reactive pattern (stride 2-5) did not well represent their true reactive muscle activity pattern. In younger adults, by contrast, the immediate response was representative, and they gradually disengaged this reactive pattern with adaptation. The fast leg’s reactive muscle activity pattern may be more difficult to capture than the slow leg because the perturbation under the fast leg (belt acceleration) more closely resembles a forward fall ^8,59,60^, which has been shown to be perceived as less severe ^59^ and to require a slower compensatory response ^61^; participants may therefore feel more comfortable exploring different strategies, producing a more variable muscle activity pattern during early adaptation. In contrast, the perturbation under the slow leg (belt deceleration) may more closely resemble a backward fall ^8,59,61^, which has been hypothesized to elicit a largely automatic response ^61^ and this response may settle into a more stereotyped pattern sooner.

In terms of visit effect, we observed a robust decrease in the recruitment of the reactive muscle activity pattern in all muscles, and an increase in context-specific recruitment in most muscles after practice. This result mirrors the behavioral finding that both age groups demonstrate robust locomotor savings (i.e., faster re-adaptation) in kinematics ^5^ and validates our conceptual framework in which two processes support locomotor adaptation: a reactive process quickly engaged for stability, and a gradually developed process reflecting the memory formation of movement patterns developed for this specific context ^7^. Here, we extend previous work by showing that the same two processes are used by both younger and older adults, and they are stored across multiple days. Specifically, the decreased reactive recruitment after practice may reflect disengagement of balance responses as the environment becomes familiar, while the increased context-specific recruitment after practice could indicate faster retrieval of the walking pattern appropriate to the current context.

Finally, the findings from EMG magnitude were largely inconsistent with both the reactive and context-specific recruitment except for the visit effect. This suggests that the decomposition and the magnitude metric capture partly different information. Specifically, overall magnitude could remain unchanged while the underlying reactive and context-specific components changed in opposing directions. The decomposition would allow us to capture this change because it reflects changes in both direction and magnitude. Conversely, an effect present in overall magnitude can be distributed across the reactive and context-specific subcomponents and therefore not reach significance in any component alone.

In summary, our results demonstrate that aging impacts the practice-induced modulation of reactive muscle recruitment but not context-specific recruitment, and that both processes are modulated during adaptation within a day and support faster re-adaptation across days. The classic control theory view of motor adaptation includes two controllers: a feedforward controller that predicts movement from the current internal model of the body and the environment, and a feedback controller that corrects for unexpected errors ^62^. Feedback-governed parameters are commonly thought to adjust immediately after an environmental transition, change little over adaptation, and show no aftereffect, whereas feedforward-governed parameters adapt slowly with time in the new environment and show aftereffects ^30,39,63,64^. Other evidence also suggests learning-dependent adjustment of the feedback control pathway ^8,10,65,66^. Here, the reactive muscle activity pattern occurred immediately after the environmental transition and continued to change during adaptation, suggesting it is generated by the feedback controller and adjusted by learning-dependent predictive gain. We therefore interpret the reactive muscle pattern as a feedback-generated response that the feedforward controller modulates during learning. In contrast, the context-specific muscle activity pattern developed gradually, so we interpret it as a feedforward update. Together, our findings suggest that aging impacts the learning-dependent modulation of the feedback control but not the development and retrieval of the feedforward model. The degraded learning-dependent modulation could reflect older adults’ resistance to update movements because of the increased uncertainty of sensory and motor signals with age as shown in previous studies ^67,68^. The preserved feedforward update of the context-specific pattern may result from an attentional compensatory strategy older adults use to support formation and retrieval of motor memories. An age-related shift toward more prefrontal-dependent, attention-demanding control of typically automatic tasks such as walking is widely reported ^69–73^ and we previously found that older adults with higher attentional control during walking exhibited faster re-adaptation of walking patterns ^5^. Therefore, future work examining changes in reactive and context-specific muscle when attentional compensation is limited, such as in individuals with cognitive impairment, could provide more insights on the mechanisms behind the preserved feedforward update. Lastly, The mapping of the reactive and context-specific processes to the feedforward and feedback control could be further clarified by future work examining the muscle activity aftereffects during tied-walking after adaptation, because aftereffects are informative about predictive adjustments and sensorimotor recalibration during learning ^44,74^. For example, if both the reactive and context-specific components are indeed modulated by a feedforward controller as we hypothesized, we should expect to see aftereffects in both.

### Limitations and Future Directions

The main limitation of the work is that we do not have a clean method to isolate a purely reactive response. The reactive response can be transient ^9^ and dissipates within a few strides, but because EMG data are noisy, particularly during perturbation onset, we averaged four strides to obtain the estimate of the reactive muscle activity pattern. The collinearity between our measured reactive and context-specific muscle activity patterns was generally low (see Method and supplementary Table S4). Nonetheless, we cannot fully rule out that our reactive muscle activity already contains some context-specific component. Future work needs to explore methods to cleanly and reliably identify a reactive muscle activity pattern to perturbed walking in muscle activity.

Lastly, the reactive and context-specific coefficients estimated by our method capture both the magnitude and the shape of muscle activity. Future studies could develop methods to decompose these coefficients to separate the magnitude and shape contributions, clarifying whether age and practice effects arise in how the muscles are activated (shape) or how much they are activated (magnitude).

### Conclusions

After practice, older adults showed limited reduction of the recruitment of the reactive muscle activity pattern than younger adults, despite comparable changes in context-specific recruitment, indicating that aging affects the practice-induced modulation of the reactive responses while leaving the formation and retrieval of context-specific motor memories intact. Both groups nonetheless showed robust reductions in reactive recruitment and increases in context-specific recruitment after practice, with no strong evidence of a group effect, suggesting that both groups shared similar muscular mechanisms for faster re-adaptation to a perturbation after practice and that the only age difference lies in the extent of reactive modulation. In sum, our results suggest that walking rehabilitation for older adults could leverage the preserved ability to form context-specific motor memories while focusing on improving reactive responses to environmental transitions during walking.

## Materials and Methods

### Participants

This study included twenty-one older (>= 65 years old) and twenty-one younger (19-40 years old) adults from a previous study ^5^. The previous study reported the kinematic results and here we present surface electromyography (EMG) data to characterize the muscle activity patterns underlying the age difference in re-adaptation when re-encountering a perturbation after practice. All participants could walk unassisted, took no medications affecting cognition, and had no major neurological conditions, severe visual impairment, or unmanaged orthopedic conditions that would interfere with walking. The Institutional Review Board at the University of Pittsburgh approved the study, and all participants gave written informed consent.

### Muscles recorded

Surface EMG data were recorded from 10 muscles per leg including five proximal muscles: gluteus medius (GLU), vastus lateralis (VL), vastus medialis (VM), semitendinosus (SEMT), and biceps femoris (BF), and five distal muscles: medial gastrocnemius (MG), lateral gastrocnemius (LG), soleus (SOL), peroneus longus (PER), and tibialis anterior (TA).

### General Experimental Protocol

The experimental protocol was described in detail in ^5^. All participants completed three visits of split-belt treadmill walking: pre-practice, practice, and post-practice (Fig. 1a), on separate days with one to two days between visits. Each visit consisted of baseline, adaptation, and washout conditions. EMG was recorded from all 20 leg muscles in every condition. Baseline was collected during both overground and treadmill walking. During overground baseline, participants walked back and forth on a 9.2m walkway at a self-selected speed for 150 strides excluding turns. A stride was defined as two consecutive heel strikes of the same leg; heel strikes were identified in the kinematic data as the times of the ankle marker’s greatest forward excursion ^68,75,76^. During treadmill baseline, participants walked on an instrumented treadmill (Bertec, Columbus, OH, United States) with both legs moving at the same speed (tied-belt) at 0.5m/s, 1.0m/s, and 0.75m/s, corresponding to slow, medium, and fast speeds, for 150 strides each. Then, during adaptation, participants walked on the treadmill with the dominant leg at 1m/s and the non-dominant leg at 0.5m/s (split-belt). Participants experienced this novel situation as a single block of 900 strides during the “pre-” and “post-practice” visits (Fig. 1a). In contrast, during the “practice” visit, participants experienced five bouts of 250 strides of split-belt walking interleaved with four bouts of 50 strides of tied-belt walking at 0.75m/s (Fig. 1a), which has been shown to accelerate locomotor savings ^4^. Next, during washout, participants walked 450 strides at their self-selected speed overground and then 150 strides on the tied-belt treadmill at 0.75m/s. The overground and tied-belt walking were included to wash out the adaptation effect from the visit.

Participants took seated resting breaks at regular intervals to limit fatigue (dashed lines in Fig. 1a). The breaks were administered at the same location with the same durations for all participants (mean ± SD break duration: 5.35 ± 0.58 mins in older adults, 5.16 ± 0.41 mins in younger adults). Participants were instructed not to move their feet during the breaks. For safety, all participants wore a harness that did not provide weight support during regular walking or standing. Participants could hold a handrail to their left whenever needed for safety, particularly during sudden speed changes such as start, stop, and speed transitions. Participants were encouraged to let go of the handrail and look straight ahead as much as possible to minimize the impact of external hand support and visual guidance on split-belt adaptation. Heart rate and blood pressure were measured, and Borg Rating of Perceived Exertion was self-reported at each break to avoid overexertion.

### EMG Data Collection and Processing

Activity from 10 muscles per leg was recorded using surface sensors (Delsys Trigno Wireless EMG system; Delsys Inc., Natick, MA, United States). Data were collected and processed following established methods ^7,8^. To ensure consistent sensor placement across visits, sensor locations were marked with skin-safe markers on the first visit to guide placement in subsequent visits whenever possible and with participant consent.

The data were streamed at 2000Hz and collected with kinematic data in Vicon Motion Systems (Oxford, United Kingdom). All data were synchronized in time using the right-belt force plate signal along the axis of gravity (Fz), which was recorded in duplicate in native Vicon and EMG system times. Then, EMG signals were high-pass filtered to remove movement artifacts and rectified. Next, we used a second-order Butterworth filter (dual-pass) with a cutoff frequency of 30Hz, resulting in 80-dB/decade attenuation and zero-lag. Lastly, the data were time-aligned to the gait cycle based on gait events determined from kinematic data ^68,75,76^. Specifically, EMG data were divided into 12 phases: two subintervals in double-support, four intervals in single-stance, two intervals in the second double-support, and four subintervals in swing. EMG amplitude was averaged within each subinterval, yielding 12 samples per stride cycle per muscle. This procedure isolated changes in muscle activity within each gait phase from any timing changes of the gait phase (for example, double support time may change during adaptation). To compare across subjects and visits, EMG activity was then amplitude-normalized to activity during fast baseline tied-belt walking (last 40 strides preceding the last 5 strides). Fast treadmill baseline was chosen as the normalization condition because it most reliably activated all muscles recorded. As a result, EMG activity is expressed in percentage units, where zero and 100% correspond to the minimum and maximum amplitude during fast treadmill baseline walking (Fig. 1b).

Next, we used a simple data-driven method to identify poor-quality recordings. Poor recordings could arise for various reasons, such as a sensor losing skin contact or a walking speed too slow to activate certain muscles (e.g., the quadriceps in some younger adults). We computed the Euclidean norm of EMG activity per stride and flagged strides falling outside three standard deviations of the mean of the distribution pooled across all muscles, visits, and participants (less than 2.22 x 10^-16^ or higher than 13.76). In the relevant epochs, 2.80% of data were excluded (supplementary Table S4). Supplementary Figure S8 presents the time course of the EMG norm for an example muscle as a visualization of the retained data. In addition, data from one younger adult’s pre-practice visit were excluded due to data collection issues.

Finally, medium-speed tied-belt walking EMG activity was removed from the EMG data (Fig. 1b), following previous convention ^7,8^, to account for intrinsic differences across participants in regular walking EMG activity. The EMG data used in the following analyses are in unbiased percentage units, where positive values represent higher-than-baseline amplitude.

### Primary EMG Outcome Measures: Reactive and Context-specific Recruitment (β)

To characterize the change in EMG activity patterns across age groups and visits, we decomposed muscle activity into two components, a reactive component associated with balance control and a context-specific component associated with a specialized movement pattern learned from practicing split-belt walking, using the following regression model:

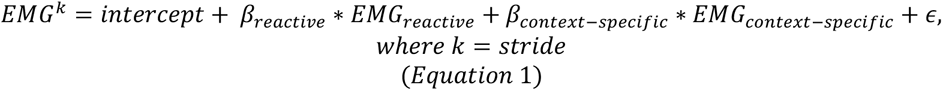

*EMG_reactive_* was measured as the initial muscle activity during early adaptation (average of the first four pre-practice strides, excluding the first transition stride from tied-belt to split-belt walking without stopping the treadmill). *EMG_context_*_–*specific*_ was measured as the late-adaptation muscle activity after practice (average of the last 40 strides of adaptation preceding the last 5 strides in the post-practice visit). In other words, *EMG_reactive_* and *EMG_context_*_–*specific*_ represent the most naïve and most practiced muscle activity during split-belt walking in our experiment. The analysis was performed at the single-subject level (see below), but a group mean visualization for the regressors (*EMG_reactive_* and *EMG_context_*_–*specific*_) is included in Fig. 1c and 1d.

The *EMG_reactive_* and *EMG_context_*_–*specific*_ muscle activity patterns were extracted for each participant and muscle. Before fitting the model, we quantified multicollinearity between the regressors using the variance inflation factor (VIF). Models were not fitted for data with high collinearity (VIF >5, ^77^), which occurred in 1.60% of the data (supplementary Table S4).

The regression coefficients *β_reactive_* and *β_context_*_–*specific*_ were estimated by ordinary least squares (OLS), solved using the normal equation in MATLAB 2024a (Mathworks, Natick, Massachusetts). For efficiency, matrix notation was used to obtain regression coefficients for all strides per muscle, visit, and participant, with a numerically stable left-division operator rather than explicit matrix inversion. Goodness of fit was measured by R^2^ and shown in supplementary Figure S7.

### General 2-level Analysis Framework

We used a two-level analysis, analogous to pipelines common in the neuroimaging literature ^78^. Specifically, we fitted a first-level linear regression model (as described above) to extract the *β_reactive_* and *β_context_*_–*specific*_ per stride. Because stride-by-stride estimates can be noisy, we averaged the outcome measures over the epoch of interest, early adaptation (strides 6 to 30) in pre- and post-practice visits. Next, we fitted a second-level linear regression model to estimate the effects of age group, visit (pre-vs post-practice), and their interaction. Specifically, for each EMG outcome measure and muscle, we fitted a linear mixed-effects model:

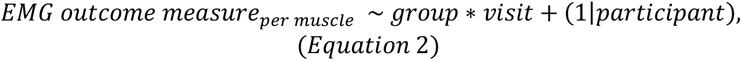

EMG outcome measures could be either *β_reactive_* or *β_context_*_–*specific*_. Group is a categorical variable with two levels (older vs younger adults), visit is a categorical variable with two levels (pre-vs post-practice), and (1|participant) represents a participant-level random intercept.

For models with a significant effect, we fitted a follow-up model to determine the specificity of the effect, i.e., whether the effect was unique to the adaptation condition and reflected practice-related muscle activity adjustments specific to this novel split-belt environment. In the follow-up model, we compared the effect observed in early adaptation (main epoch of interest) against leg-speed-specific baseline conditions by testing for a condition interaction. Specifically, we fitted:

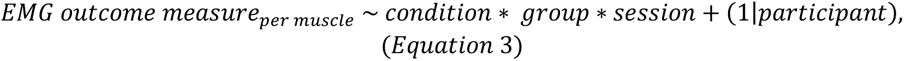

Condition is a categorical variable with two levels: early adaptation or speed-specific baseline. The speed-specific baseline was fast tied-belt walking for fast-leg adaptation data and slow tied-belt walking for slow-leg adaptation data. This comparison isolated group and visit effects unique to learning the split-belt situation (controlling two legs that move at different speeds simultaneously) from effects due simply to each leg’s walking speed. A significant interaction with a greater effect during adaptation than during speed-specific baseline would indicate that the effect observed in Equation 2 (main model) was specific to the adaptation condition.

### Secondary EMG Outcome Measures: EMG Magnitude

To compare our results with the conventional metric, we also repeated the linear mixed-effects (second-level) model to estimate the effects of age group, visit, and their interaction on EMG magnitude. EMG magnitude was measured as the Euclidean norm of the time-aligned, binned, and amplitude-normalized EMG across the 12 subintervals of a gait cycle. A higher value represents more overall muscle activity in a stride, without information about the activity pattern.

### Epoch of Interest

The main epoch of interest was early adaptation, defined as the average EMG outcome measures over strides 6 to 30 from the onset of split-belt walking. The epoch was chosen to capture the fast-fleeting reactive response to split-belt perturbation. Analysis of the young adults’ pre-practice data, the dataset most similar to a stereotypical motor-adaptation protocol, suggested that by focusing on strides 6 to 30, the window immediately following *EMG_reactive_*, we could capture the dynamics of the reactive process in at least 85% of the data, measured by percentage of participants whose *β_reactive_* does not reach a plateau ^79^ within the first 30 strides (data not shown).

### Statistical Analysis

The primary analysis was to identify the interaction effect of age group and visit on the EMG outcome measures: *β_reactive_* and *β_context_*_–*specific*_. We extracted the interaction effect size, quantified by 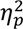, and p-values per outcome measure per muscle using the second-level linear mixed-effects model described above. Because violations of normality (checked by Shapiro-Wilk test ^80^) and homoscedasticity (checked by Bartlett’s test ^81^) were common, non-parametric tests were performed. Confidence intervals for all model coefficients and effect sizes were estimated via a cluster bootstrap (5,000 iterations, with subjects as the resampling unit stratified by group ^82^). Permutation testing with 5,000 iterations was performed to extract the p-values using aovperm (permuco package) in R (version 4.5.1). Multiple-comparison corrections (for 20 muscles) were performed using the Benjamini-Hochberg procedure to control the false discovery rate (FDR ^31^) at 0.05. We reported the observed model coefficient for the interaction term, 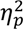 (estimated from lme4 and effectsize package) with bootstrapped 95% confidence interval, and the adjusted permutation p-values.

The secondary analysis was to identify the effects of age group and visit. The same analysis described above for the interaction effect was performed. Age group and visit effects were treated as two separate families, so the FDR procedure corrected 20 comparisons each.

When a significant interaction or main effect was found, a follow-up specificity analysis was performed to determine whether the observed effect was specific to the early adaptation epoch. A linear mixed-effects model with condition interaction was fitted as shown in Equation 3 using data from early adaptation stacked with the leg-speed-specific baseline condition. If the group by visit interaction was specific to the adaptation condition, we expected a significant condition by group by visit interaction in the follow-up model and the magnitude of the group by visit coefficient in adaptation should be larger than that in baseline. Therefore, a one-tailed comparison of |*β*_7*roup*∗*session*_ adaptation| − |*β*_7*roup*∗*session*_ baseline| was tested by permutation testing via the Freedman-Lane algorithm (5,000 permutations). Briefly, the term of interest was permuted by reshuffling the subject-level residuals from the corresponding reduced model. The EMG outcome measures in the speed-specific baseline condition are included in supplementary Figure S3 and S4 to visually compare the effect in the adaptation condition (supplementary Figure S1 and S2) against the speed-specific baseline. No multiple-comparison correction was performed in the follow-up analysis.

To explore the consistency of our findings against a conventional metric, all analyses were repeated for EMG magnitude.

A significance level of *α* = 0.05 was used for all analyses. Data processing, outcome extraction, and first-level models were fitted in MATLAB. Second-level linear mixed-effects models and specificity analyses were performed in R (version 4.5.1).

## Acknowledgements

The authors thank Dr. Krista Fjeld, Jiwon Choi, Elan Albalak, Shaina Patel, Abby Somich, Emma Stein, Shaoyi Li, and Devin Stafford for their tremendous help during data collection. The authors thank Dr. Marcela Gonzalez-Rubio for the insightful feedback on the analysis framework, the manuscript, and result presentation, and help during data collection. The authors thank Dr. Chase Rock and Jiwon Choi for the insights on data analysis and feedback on the manuscript. The authors thank Nathan Brantly and Adwoa A. Awuah for their feedback on the manuscript. The authors thank Dr. Dulce M. Mariscal and Dr. Pablo A. Iturralde for inspiration on the EMG model to characterize the two processes underlying locomotor adaptation.

## Author contributions

S.L. and G.T. conceptualized and designed research; A.W. and G.T. advised on data analysis and interpretation and reviewed and edited the manuscript; G.T. acquired funding; S.L. performed research, analyzed data, and wrote the first draft of the paper.

## Data availability statement

All data and code needed to reproduce the results in the manuscript will be publicly available on open science framework (OSF) and GitHub upon publication of the manuscript.

## Additional Information

### Disclosure

None

### Declaration of AI Usage

Claude AI was used to develop the data analysis code in R and result visualization code in MATLAB. All code was verified by SL. Claude AI was also used to refine the language for grammatically errors, consistency, and clarity. All content was generated originally.

