## Supplementary Materials for "Aging limits the reduction of reactive muscle activity to perturbations but spares the retrieval of context-specific patterns with practice"

**Corresponding Author:** Dr. Gelsy Torres-Oviedo

### Figures

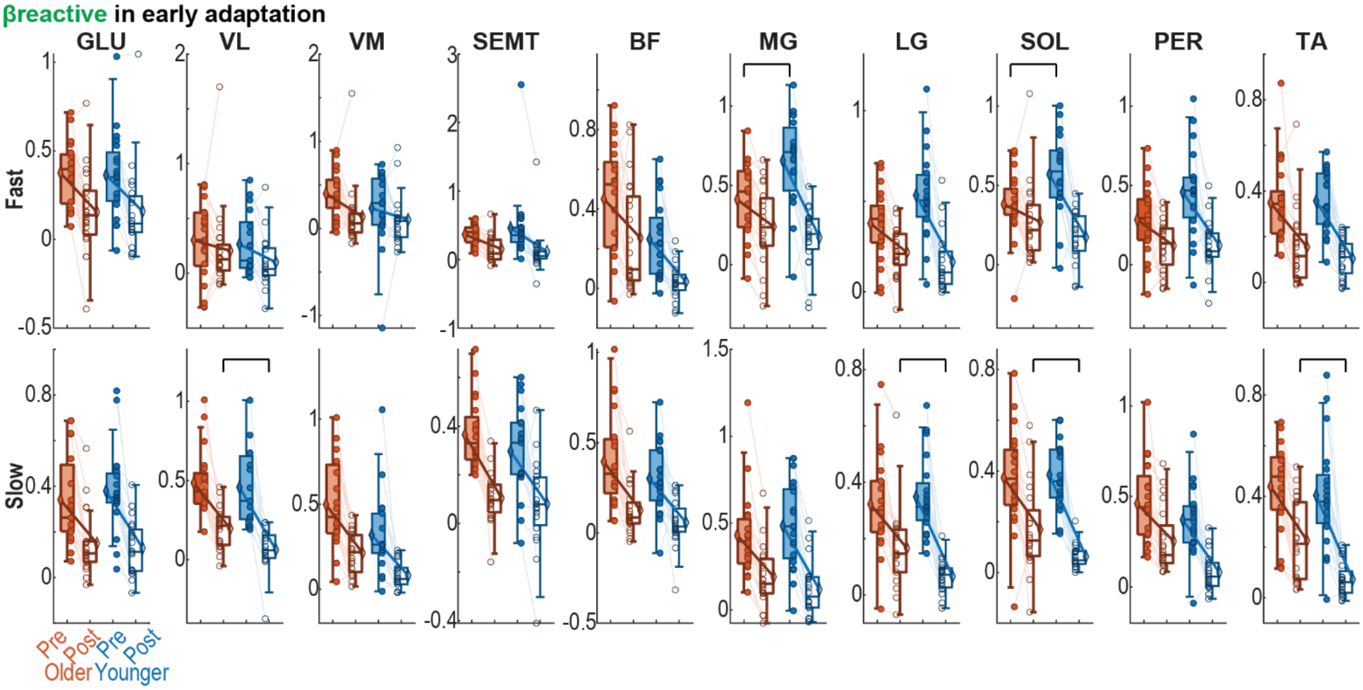


**Figure S1. Linear mixed-effect model $\beta_{reactive}$ data in all muscles.** Box plot visualization of the $\beta_{reactive}$ data used in the second-level linear mixed-effect model testing the effects of group, visit, and their interaction. Fast leg muscles are shown in top row from proximal to distal. Slow leg muscles are shown in bottom row. *Each dot represents one participant (orange, older adults; blue, younger adults; filled, pre-practice; open, post-practice). Thin lines connect data from the same participant. Group means are shown as diamonds connected by thick lines. In muscles that showed a group-by-visit interaction with medium or large effect size, post-hoc comparisons between group within each visit were performed. Significant post-hoc comparisons after Benjamini–Hochberg false discovery rate (FDR)*[1] *correction of 40 comparisons are shown with a solid line bracket.*

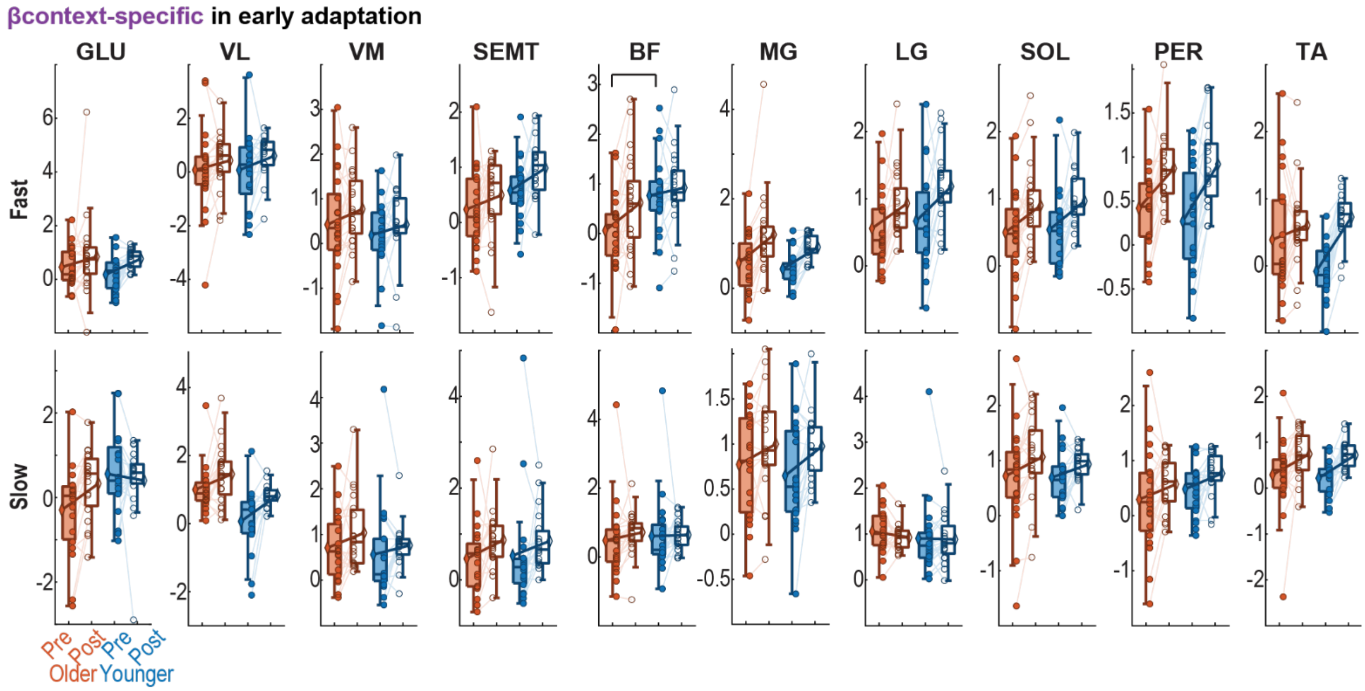


**Figure S2. Linear mixed-effect model $\boldsymbol{\beta_{context-specific}}$ data in all muscles.** Box plot visualization of the $\boldsymbol{\beta_{context-specific}}$ data used in the second-level linear mixed-effect model testing the effects of group, visit, and their interaction. Fast leg muscles are shown in top row from proximal to distal. Slow leg muscles are shown in bottom row. *Each dot represents one participant (orange, older adults; blue, younger adults; filled, pre-practice; open, post-practice). Thin lines connect data from the same participant. Group means are shown as diamonds connected by thick lines. In muscles that showed a group-by-visit interaction with medium or large effect size, post-hoc comparisons between group within each visit were performed. Significant post-hoc comparisons after Benjamini–Hochberg false discovery rate (FDR)* [1] *correction of 40 comparisons are shown with a solid line bracket.*

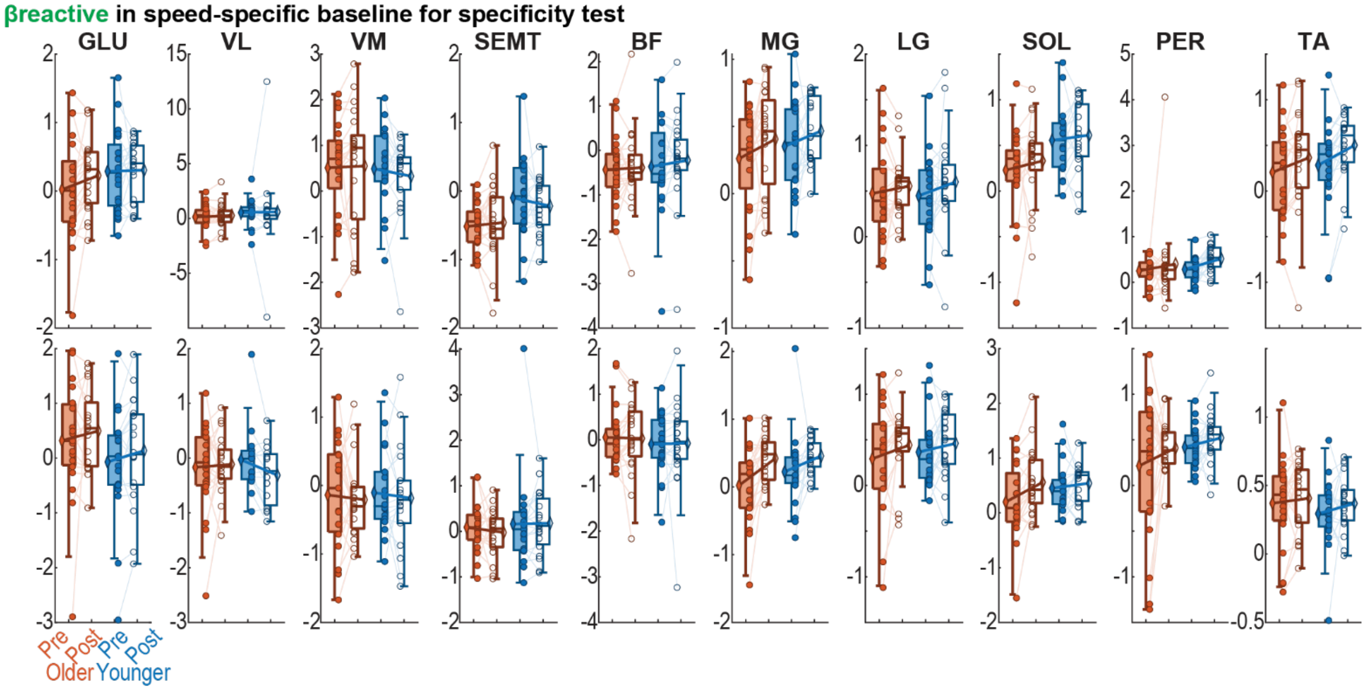

**Figure S3. $\boldsymbol{\beta_{reactive}}$ data in the speed-specific baseline condition used in the follow-up** specificity **analysis.** *Box plot visualization of the $\beta_{reactive}$ data used in the follow-up specificity analysis testing whether the effect observed in Figure S1 is specific to the early adaptation condition. If it is specific, we expect either no effects or smaller effects than the ones observed in the early adaptation condition in Figure S1. Fast leg muscles are shown in top row from proximal to distal. Slow leg muscles are shown in bottom row. Each dot represents one participant (orange, older adults; blue, younger adults; filled, pre-practice; open, post-practice). Thin lines connect data from the same participant. Group means are shown as diamonds connected by thick lines.*

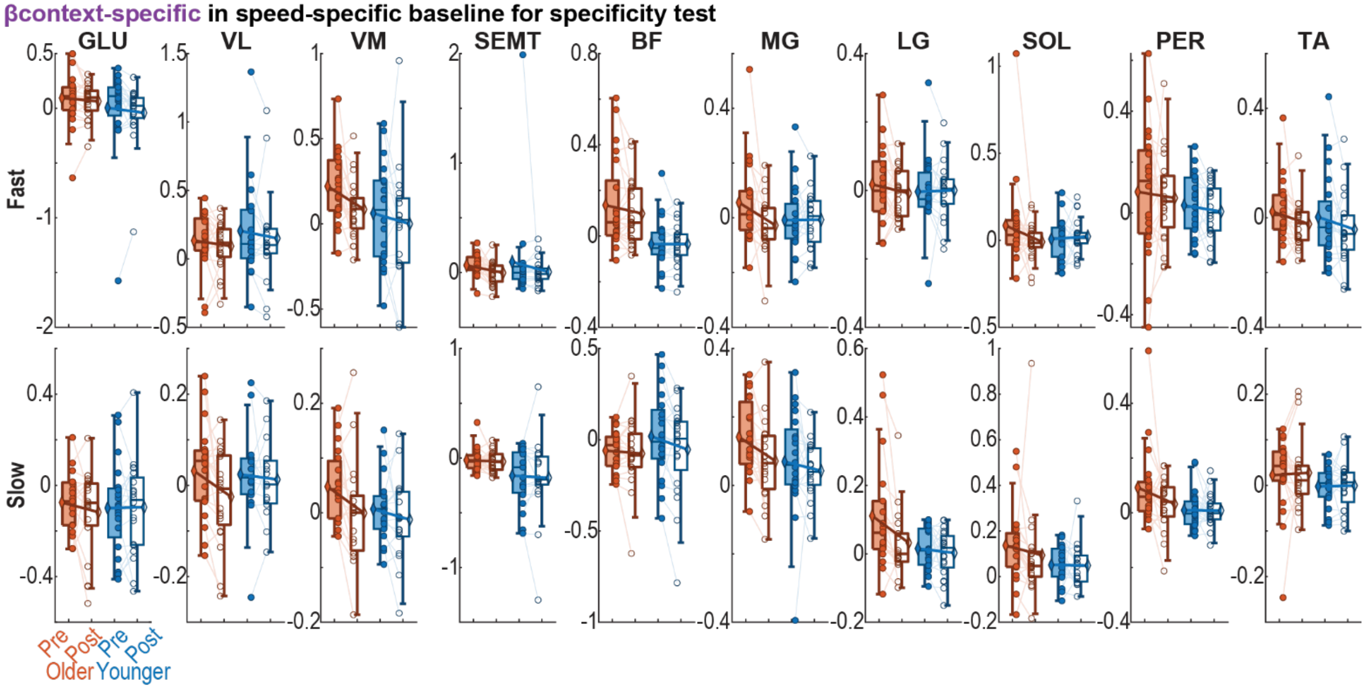


**Figure S4. $\boldsymbol{\beta_{context-specific}}$ data in the speed-specific baseline condition used in the follow-up** specificity **analysis.** *Box plot visualization of the $\beta_{context-specific}$ data used in the follow-up specificity analysis testing whether the effect observed in Figure S1 is specific to the early adaptation condition. If it is specific, we expect either no effects or smaller effects than the ones observed in the early adaptation condition in Figure S1. Fast leg muscles are shown in top row from proximal to distal. Slow leg muscles are shown in bottom row. Each dot represents one participant (orange, older adults; blue, younger adults; filled, pre-practice; open, post-practice). Thin lines connect data from the same participant. Group means are shown as diamonds connected by thick lines.*

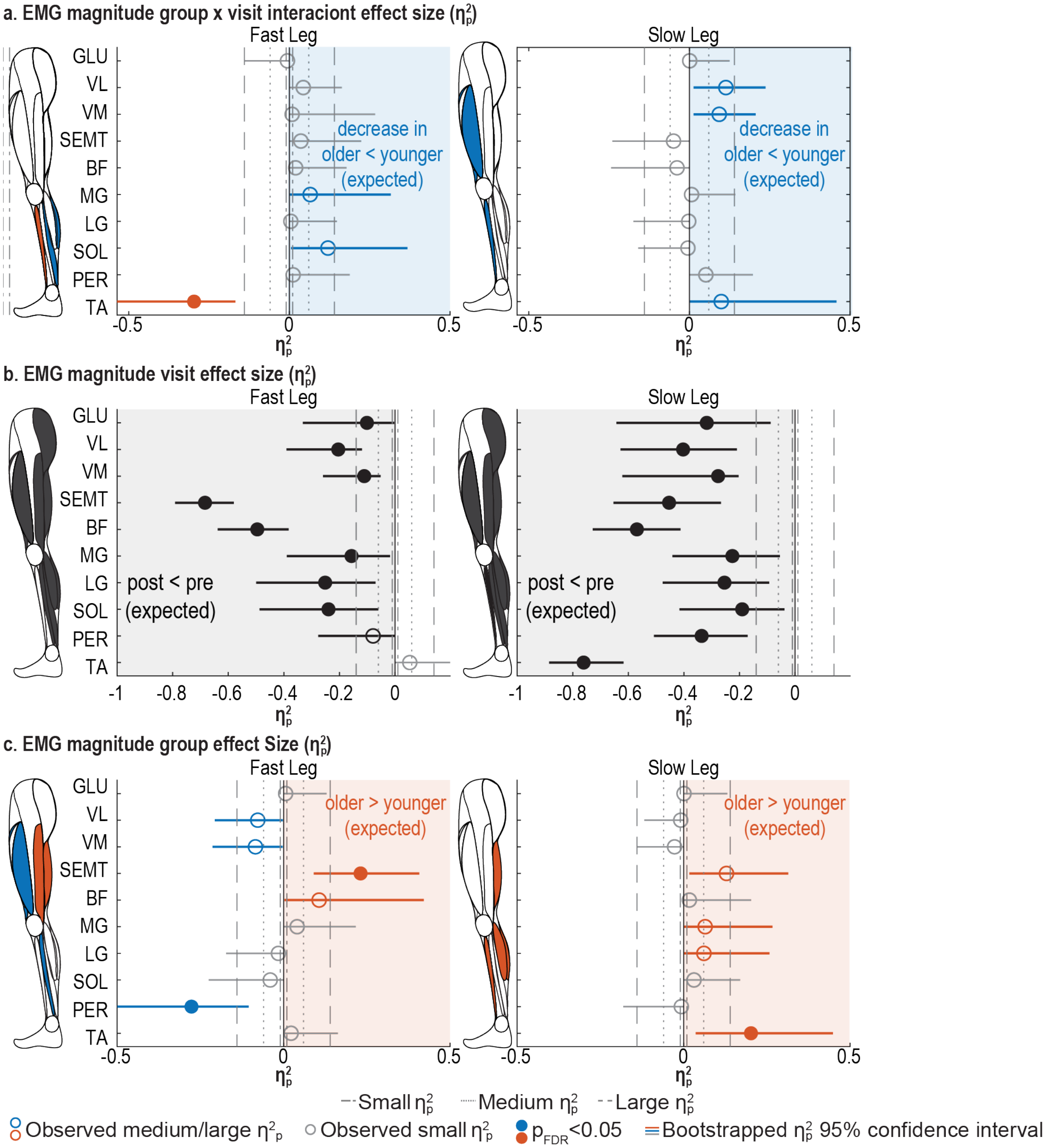


***Figure S5. Age-by-visit interaction (a), main effects of visit (c) and group (e) in EMG magnitude. (a)*** *Overview of the interaction effect size $\eta_p^2$ for EMG magnitude. Each dot represents the observed effect size; the line represents the 95% confidence interval from bootstrapping. Filled dots indicate a significant interaction after Benjamini–Hochberg false discovery rate (FDR) [1] correction of 40 comparisons are shown with a solid line bracket.. Directions of the expected effects are shown with colored background. In panel a, the expected direction is marked in blue where older adults change less than younger adults. Medium or large effects in the expected direction are colored in blue and in orange in the unexpected direction; small or negligible effects are grey. Dot-dashed, dotted, and dashed lines mark thresholds for small (0.01), medium (0.06), and large (≥0.14) effect sizes [2].* ***(b)*** *Overview of the visit effect $\eta_p^2$ for EMG magnitude. The grey background marks the expected direction where the EMG magnitude decreased after practice. Medium or large effects are colored black in the expected direction; small or negligible effects are grey.* ***(c)*** *Overview of the group effect size $\eta_p^2$ for EMG magnitude. The orange background marks the expected*

*direction with higher EMG magnitude in older than younger adults. Medium or large effects are colored orange in the expected direction and blue in the unexpected direction; small or negligible effects are grey.*

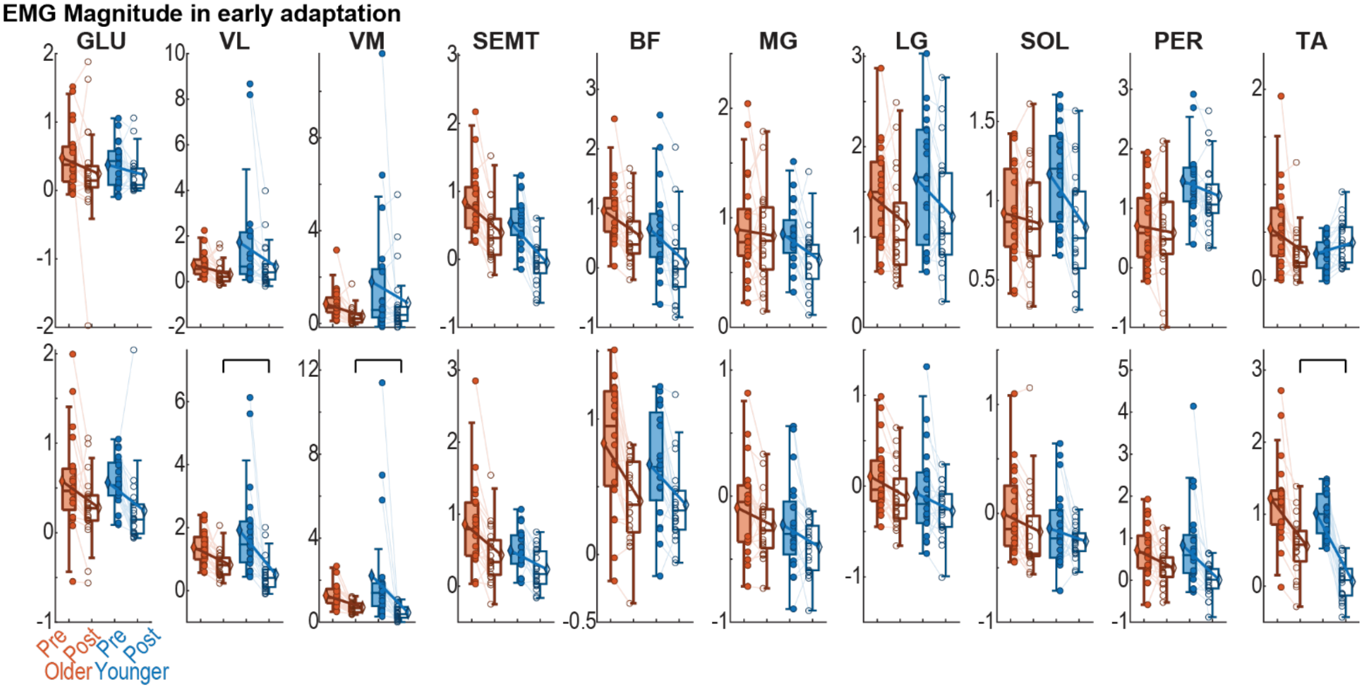


**Figure S6. Linear mixed-effect model data of EMG magnitude in all muscles.** *Box plot visualization of the EMG magnitude data used in the second-level linear mixed-effect model testing the effects of group, visit, and their interaction. Fast leg muscles are shown in top row from proximal to distal. Slow leg muscles are shown in bottom row. Each dot represents one participant (orange, older adults; blue, younger adults; filled, pre-practice; open, post-practice). Thin lines connect data from the same participant. Group means are shown as diamonds connected by thick lines. In muscles that showed a group-by-visit interaction with medium or large effect size, post-hoc comparisons between group within each visit were performed. Significant post-hoc comparisons after Benjamini–Hochberg false discovery rate (FDR)* [1] *correction of 40 comparisons are shown with a solid line bracket.*

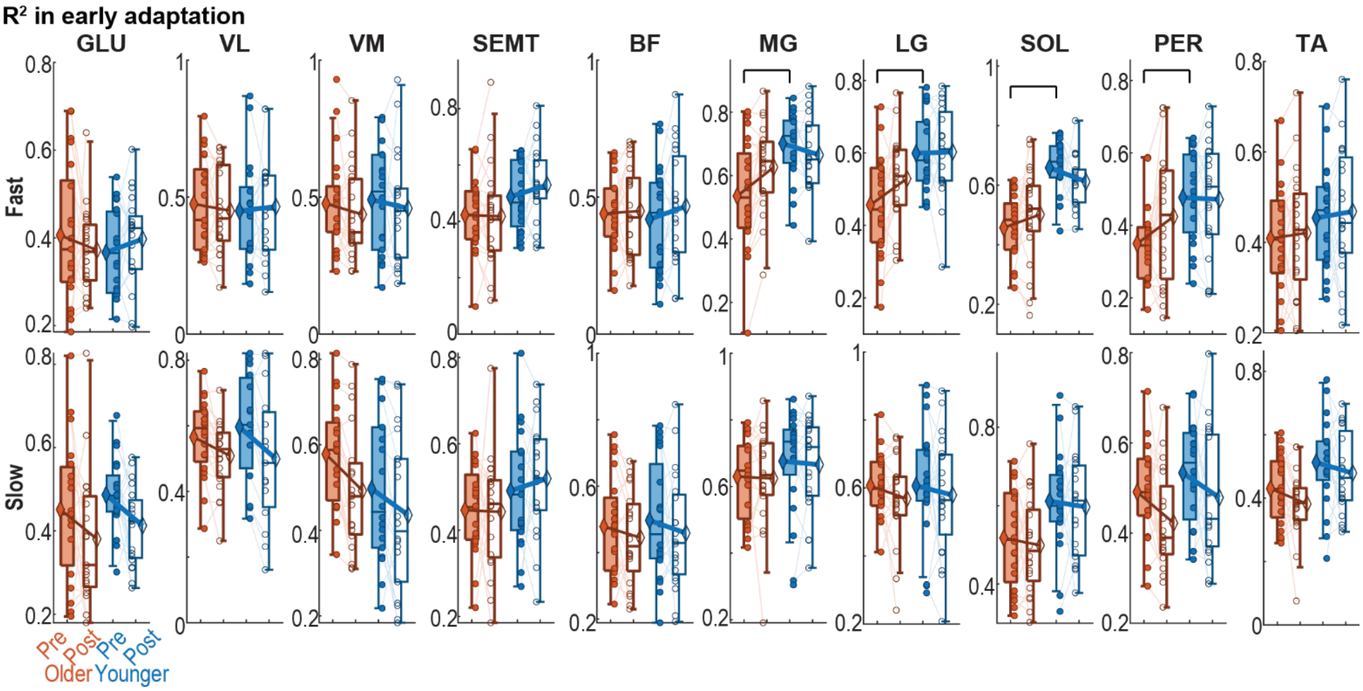


**Figure S7. Goodness of fit ($R^2$) of the first-level model used to extract the $\beta_{reactive}$ and $\beta_{context-specific}$.** *The first-level model (main text Figure 1b) was fitted separately for each participant and muscle; the resulting $R^2$ is shown with fast leg muscles on the top row from proximal to distal and slow leg muscles on the bottom row. Each dot represents one participant (orange, older adults; blue, younger adults; filled, pre-practice; open, post-practice). Thin lines connect data from the same participant. Group means are shown as diamonds connected by thick lines. In muscles that showed a group-by-visit interaction with medium or large effect size, post-hoc comparisons between group within each visit were performed. Significant post-hoc comparisons after Benjamini–Hochberg false discovery rate (FDR)* [1] *correction of 40 comparisons are shown with a solid line bracket.*

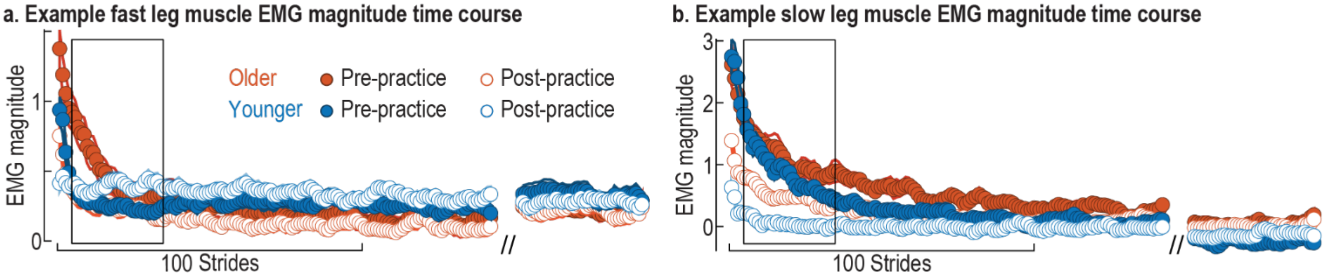


**Figure S8. Time course of EMG magnitude in example fast (a) and slow (b) leg muscles (*tibialis anterior, TA*) *during early and late adaptation.*** *In both panels, older adults are shown in orange, younger adults in blue; pre-practice and post-practice data are in closed and open circles, respectively. Each dot represents the average of 5 consecutive strides, and shaded area represents the standard error. Except for fast-leg TA in younger adults, EMG magnitude decreased during adaptation in both groups and was smaller at the post-practice.*

### Tables

**Table S1. $\beta_{reactive}$ and $\beta_{context-specific}$ follow-up specificity test result p-value.**
Specificity tests evaluated whether observed effects were unique to the adaptation condition with effect size larger than baseline. Analyses were restricted to muscles exhibiting a significant effect in the interaction, group, or visit effects reported above and the symbol "/" indicates muscles where the specificity analysis was not performed. All p-values are uncorrected for multiple comparisons because the specificity test is conditional on significant p-values from previous results that have been corrected. Specificity tests that are significant are formatted in bold.

| | Muscle | $\beta_{reactive}$ Follow-up p-values | | | $\beta_{context-specific}$ Follow-up p-values | | |
|---|---|---|---|---|---|---|---|
| | | Interaction Effect | Visit Effect | Group Effect | Interaction Effect | Visit Effect | Group Effect |
| FAST | GLU | / | **0.004** | / | / | **0.004** | / |
| | VL | / | 0.1 | / | / | 0.05 | / |
| | VM | / | 0.24 | / | / | / | / |
| | SEMT | / | **<.001** | / | / | 0.07 | / |
| | BF | / | **<.001** | 0.33 | / | 0.5 | / |
| | MG | **0.01** | **<.001** | / | / | **0.002** | / |
| | LG | **0.01** | **<.001** | / | / | **0.03** | / |
| | SOL | **0.05** | **<.001** | / | / | **0.007** | / |
| | PER | / | **<.001** | / | / | **<.001** | / |
| | TA | / | **<.001** | / | **0.002** | **<.001** | / |
| SLOW | GLU | / | **<.001** | / | / | / | / |
| | VL | / | **<.001** | / | / | 0.11 | **0.03** |
| | VM | / | **<.001** | 0.06 | / | / | / |
| | SEMT | / | **0.009** | / | / | **0.03** | / |
| | BF | / | **0.003** | / | / | / | / |
| | MG | / | **<.001** | / | / | 0.18 | / |
| | LG | 0.10 | **<.001** | / | / | / | / |
| | SOL | / | **<.001** | / | / | 0.12 | / |
| | PER | / | **<.001** | 0.47 | / | 0.06 | / |
| | TA | / | **<.001** | / | / | **<.001** | / |

**Table S2. EMG magnitude linear mixed effect model results.** *The reported $\beta$ and $\eta_p^2$ are the model estimate derived from the observed data. The 95% confidence interval was obtained via bootstrapping. $P$-values were computed by permutation testing and adjusted for multiple comparisons across 20 muscles using the Benjamini–Hochberg false discovery rate (FDR) procedure [1]. Effects remaining significant after correction are formatted in bold. Consistent with the color schemes in the Figure S5, observed medium or large interaction effect sizes ($\eta_p^2$ >= 0.06 [2]) are shown in orange when the change in younger adults exceeds that in older adults (matching expectation), and in blue when the change in older adults exceeds that in younger adults (contrary to expectation). Small or negligible effects are in grey. Observed medium or large visit effect in the expected direction are shown in black. Observed medium to large visit effect in the opposite direction where EMG magnitude increased after practice is shown in yellow. Observed medium or large group effect are shown in blue when EMG magnitude is greater in older adults and in orange when the magnitude is greater in younger adults. We expected older adults to recruit have larger EMG magnitude.*

| | Muscle | $\beta_{interaction}$ with bootstrapped 95% CI | $\eta_p^2$ | p | $\beta_{visit}$ with bootstrapped 95% CI | $\eta_p^2$ | p | $\beta_{group}$ with bootstrapped 95% CI | $\eta_p^2$ | p |
|---|---|---|---|---|---|---|---|---|---|---|
| FAST | GLU | -0.09 [-0.44, 0.24] | 0.01 | 0.76 | **-0.14 [-0.31, 0.02]** | **0.1** | **0.04** | 0.11 [-0.13, 0.35] | 0.01 | 0.67 |
| | VL | 0.61 [-0.20, 1.64] | 0.04 | 0.36 | **-1.03 [-2.06, -0.28]** | **0.2** | **0.003** | -0.96 [-2.19, 0.02] | 0.08 | 0.19 |
| | VM | 0.38 [-0.69, 1.75] | 0.01 | 0.76 | **-0.91 [-2.29, 0.11]** | **0.11** | **0.03** | -0.94 [-2.31, 0.16] | 0.08 | 0.19 |
| | SEMT | 0.14 [-0.08, 0.35] | 0.04 | 0.39 | **-0.59 [-0.75, -0.42]** | **0.68** | **<0.001** | **0.31 [0.06, 0.58]** | **0.23** | **0.01** |

| | | | | | | | | | | |
|---|---|---|---|---|---|---|---|---|---|---|
| | BF | 0.14 [-0.17, 0.45] | 0.02 | 0.59 | **-0.56 [-0.80, -0.35]** | **0.5** | **<0.001** | 0.31 [-0.08, 0.68] | 0.11 | 0.14 |
| | MG | 0.18 [-0.02, 0.38] | 0.06 | 0.33 | **-0.24 [-0.35, -0.11]** | **0.16** | **0.01** | 0.04 [-0.18, 0.28] | 0.04 | 0.37 |
| | LG | 0.08 [-0.31, 0.47] | 0.004 | 0.76 | **-0.41 [-0.72, -0.09]** | **0.25** | **0.001** | -0.17 [-0.55, 0.22] | 0.02 | 0.52 |
| | SOL | 0.26 [0.05, 0.47] | 0.12 | 0.22 | **-0.33 [-0.50, -0.16]** | **0.24** | **0.002** | -0.24 [-0.42, -0.05] | 0.04 | 0.37 |
| | PER | 0.14 [-0.27, 0.50] | 0.01 | 0.75 | -0.25 [-0.49, 0.002] | 0.08 | 0.07 | **-0.75 [-1.12, -0.38]** | **0.28** | **0.007** |
| | TA | **-0.39 [-0.58, -0.23]** | **0.3** | **0.005** | 0.12 [0.06, 0.19] | 0.05 | 0.13 | 0.27 [0.09, 0.48] | 0.02 | 0.52 |
| SLOW | GLU | 0.02 [-0.26, 0.29] | 0.001 | 0.88 | **-0.32 [-0.49, -0.12]** | **0.32** | **<0.001** | 0.01 [-0.25, 0.29] | 0.001 | 0.83 |
| | VL | 0.86 [0.17, 1.68] | 0.11 | 0.22 | **-1.42 [-2.21, -0.77]** | **0.4** | **<0.001** | -0.55 [-1.34, 0.12] | 0.01 | 0.66 |
| | VM | 1.20 [0.17, 2.45] | 0.09 | 0.22 | **-1.76 [-3.02, -0.75]** | **0.28** | **<0.001** | -0.94 [-2.25, 0.12] | 0.03 | 0.44 |
| | SEMT | -0.17 [-0.40, 0.06] | 0.05 | 0.36 | **-0.25 [-0.41, -0.10]** | **0.45** | **<0.001** | 0.37 [0.09, 0.69] | 0.13 | 0.1 |
| | BF | -0.12 [-0.31, 0.08] | 0.04 | 0.39 | **-0.29 [-0.45, -0.15]** | **0.57** | **<0.001** | 0.15 [-0.12, 0.41] | 0.02 | 0.52 |
| | MG | 0.05 [-0.13, 0.23] | 0.01 | 0.81 | **-0.17 [-0.31, -0.05]** | **0.23** | **0.002** | 0.12 [-0.11, 0.35] | 0.06 | 0.22 |
| | LG | -0.03 [-0.25, 0.20] | 0.002 | 0.87 | **-0.19 [-0.41, 0.001]** | **0.25** | **0.003** | 0.19 [-0.10, 0.47] | 0.06 | 0.27 |
| | SOL | -0.04 [-0.21, 0.14] | 0.005 | 0.76 | **-0.12 [-0.27, 0.03]** | **0.19** | **0.004** | 0.13 [-0.09, 0.35] | 0.03 | 0.41 |
| | PER | 0.40 [-0.10, 0.96] | 0.05 | 0.36 | **-0.82 [-1.31, -0.41]** | **0.34** | **<0.001** | -0.12 [-0.67, 0.38] | 0.01 | 0.66 |
| | TA | 0.30 [0.004, 0.57] | 0.1 | 0.22 | **-0.95 [-1.11, -0.81]** | **0.76** | **<0.001** | **0.19 [-0.09, 0.49]** | **0.2** | **0.02** |

**Table S3. EMG magnitude specificity test result p-value.**

Specificity tests evaluated whether observed effects were unique to the adaptation condition with effect size larger than baseline. Analyses were restricted to muscles exhibiting a significant effect in the interaction, group, or visit effects reported above and the symbol "/" indicates muscles where the specificity analysis was not performed. All p-values are uncorrected for multiple comparisons because the specificity test is conditional on significant p-values from previous results that have been corrected. Specificity tests that are significant are formatted in bold.

| | | EMG Magnitude Follow-up p-value | | |
|---|---|---|---|---|
| **Muscle** | | **Interaction Effect** | **Visit Effect** | **Group Effect** |
| FAST | GLU | / | / | / |
| | VL | / | **0.005** | / |
| | VM | / | / | / |
| | SEMT | / | **<0.001** | / |
| | BF | / | **<0.001** | / |
| | MG | / | **0.003** | / |
| | LG | / | **0.02** | / |
| | SOL | / | **0.02** | / |
| | PER | / | / | **0.004** |
| | TA | **<0.001** | / | / |
| SLOW | GLU | / | **0.01** | / |
| | VL | / | **0.01** | / |
| | VM | / | **<0.001** | / |
| | SEMT | / | **0.004** | / |
| | BF | / | **0.02** | / |
| | MG | / | / | / |
| | LG | / | / | / |
| | SOL | / | / | / |

| | | | |
|---|---|---|---|
| PER | / | **0.001** | / |
| TA | / | **<0.001** | / |

**Table S4. Summary of participant exclusions for linear mixed-effects model fitting.**
Number of participants excluded from linear mixed-effects model fitting due to poor electromyography (EMG) signal quality or high multicollinearity, defined as a variance inflation factor (VIF) greater than 5. Values represent counts of excluded individuals per age and visit combination. Because VIF values were identical across pre- and post-practice sessions, they are reported once per age group. For reference, total number of participants with available EMG data is 20 for younger adults pre-practice visit due to issues in data collection, and 21 in all other cases.

| **Muscle** | | **Bad Recording Quality** | | **High VIF** | |
|---|---|---|---|---|---|
| | | **Older** | **Younger** | **Older** | **Younger** |
| FAST | GLU | 0 | 0 | 0 | 0 |
| | VL | 0 | 3 | 0 | 0 |
| | VM | 0 | 1 | 0 | 1 |
| | SEMT | 0 | 0 | 0 | 0 |
| | BF | 0 | 0 | 0 | 0 |
| | MG | 0 | 0 | 1 | 1 |
| | LG | 0 | 0 | 0 | 1 |
| | SOL | 0 | 0 | 0 | 2 |
| | PER | 0 | 0 | 1 | 1 |
| | TA | 0 | 0 | 0 | 0 |
| SLOW | GLU | 1 | 0 | 1 | 0 |
| | VL | 1 | 5 | 0 | 0 |
| | VM | 1 | 2 | 1 | 0 |
| | SEMT | 1 | 0 | 0 | 1 |
| | BF | 1 | 0 | 0 | 0 |
| | MG | 2 | 0 | 0 | 0 |
| | LG | 2 | 0 | 1 | 0 |
| | SOL | 1 | 0 | 0 | 1 |
| | PER | 1 | 0 | 0 | 0 |
| | TA | 1 | 0 | 0 | 0 |
